# Task-dependent mixed selectivity in the subiculum

**DOI:** 10.1101/2020.06.06.129221

**Authors:** Debora Ledergerber, Claudia Battistin, Jan Sigurd Blackstad, Richard J. Gardner, Menno P. Witter, May-Britt Moser, Yasser Roudi, Edvard I. Moser

## Abstract

CA1 and subiculum (SUB) connect the hippocampus to numerous output regions. Cells in both areas have place-specific firing fields, although they are more dispersed in SUB. Weak responses to head direction and running speed have been reported in both regions. However, how such information is encoded in CA1 and SUB, and the resulting impact on downstream targets, is poorly understood. Here we estimate the tuning of simultaneously recorded CA1 and SUB cells to position, head direction, and speed. Individual neurons respond conjunctively to these covariates in both regions but the degree of mixed representation is stronger in SUB, and more so during goal-directed spatial navigation than free foraging. Each navigational variable could be decoded with higher precision, from a similar number of neurons, in SUB than CA1. The findings point to a possible contribution of mixed-selective coding in SUB to efficient transmission of hippocampal representations to widespread brain regions.

## Introduction

The hippocampus has a well-established role in mnemonic and navigational functions of the brain (Hasselmo, 2012; Morris, 2007; O’Keefe and Nadel, 1978; Scoville and Milner, 1957; Squire, 1992). Its interplay with other cortical regions is thought to be indispensable for the formation and retrieval of episodic and positional memories (Buzsáki, 1989; McClelland et al., 1995; Squire, 1992; Squire et al., 2015; Winocur and Moscovitch, 2011). The subiculum (SUB) has an important anatomical position as an interface between the hippocampus and other brain areas (Cappaert et al., 2015; O’Mara, 2006). It is the major long-range projection area of the hippocampus, the origin of a substantial part of the fornix, and the source of large parts of non-fornical output reaching a range of cortical and subcortical downstream areas. Along with the CA1 area, it sends outputs to infralimbic, prefrontal, orbitofrontal and medial and lateral entorhinal cortices (MEC and LEC) and to subcortical structures including the septal complex, the mammillary nucleus, the hypothalamus, the thalamus and the amygdala (Cappaert et al., 2015; Cembrowski et al., 2018a; Ishizuka, 2001; O’Mara et al., 2001). However, even though CA1 and SUB have many overlapping target areas, their patterns of connectivity are very different: the majority of CA1 neurons each send collateral projections to at least two targets elsewhere in the brain, whereas in case of the subiculum such branching is only observed in a minority of neurons (Bienkowski et al., 2018; Cembrowski et al., 2018b; Naber and Witter, 1998; Witter, 2006). This difference raises the possibility that the hippocampus uses CA1 and SUB outputs differentially to distribute information to downstream brain regions.

While neuronal representations in CA1 have been known for half a century to show strong spatial selectivity, as expressed in place cells (O’Keefe and Dostrovsky, 1971; O’Keefe and Nadel, 1978), data from SUB is scarce and it has remained elusive what SUB adds to the hippocampal computation. Subpopulations of SUB neurons have broad spatial firing fields (Sharp, 1997, 2006; Sharp and Green, 1994), some of which, known as boundary-vector cells, are oriented in parallel to elongated geometric boundaries (Lever et al., 2009; Stewart et al., 2014). Other SUB cells have been shown to encode the animal’s axis of movement (heading direction) when animals navigate on elevated multidirectional tracks (Olson et al., 2016). In all tasks where SUB neurons have been recorded, they appear to be more broadly tuned to features of behavior or environment than neurons in other regions of the hippocampal formation. SUB is likely to receive navigational input from narrowly tuned place cells in CA1 (O’Keefe, 1976; O’Keefe and Dostrovsky, 1971), from grid cells in medial entorhinal cortex (MEC) and pre- and parasubiculum (Boccara et al., 2010; Fyhn et al., 2004; Hafting et al., 2005), from head direction cells (Taube and Burton, 1995; Taube et al., 1990), border cells and object vector cells (Høydal et al., 2019; Solstad et al., 2008) in the same regions, and from speed cells in the hippocampus and MEC (Kropff et al., 2015). The specificity of these putative inputs brings up the question of what the broader representations in SUB add to the output of the hippocampal formation.

Navigation-tuned cells in the hippocampal formation express similar types of information in multiple environments. This is in contrast to neuronal representations in many other brain areas, where spatial selectivity is apparent only under task conditions relevant to the brain region, such as in prefrontal cortex (Jung et al., 1998; Padilla-Coreano et al., 2019; Pratt and Mizumori, 2001), posterior parietal cortex (Nitz, 2006; Whitlock et al., 2012), primary visual cortex (Goltstein et al., 2018; Saleem et al., 2018), amygdala (Peck et al., 2014) and nucleus accumbens (Lansink et al., 2012; Mulder et al., 2005). Navigational information in these regions may be derived from representations in the hippocampus (Remondes and Wilson, 2013; Spellman et al., 2015), but it remains unclear whether, and how, outputs from hippocampus would be modified in a task-specific manner before reaching these diverse regions. Given the potential role of SUB in distributing hippocampal output to widespread regions of the brain (Cappaert et al., 2015; Gigg, 2006; O’Mara, 2006), we hypothesized that, rather than generating *de novo* representations, SUB modifies representations from upstream neural populations in CA1, presubiculum and entorhinal cortex to facilitate decoding by downstream regions during hippocampal-dependent behaviors.

It has been suggested that networks consisting of neurons that encode multiple stimulus features simultaneously or conjunctively, using a ‘mixed selectivity’ code, have several computational advantages over networks where neurons respond predominantly to single features (Miller et al., 1996; Rigotti et al., 2013). Besides a high representational capacity, networks with high-dimensional coding in individual neurons have the advantage that a wide span of task-relevant aspects is accessible to linear classifiers, as the number of classifications that can be performed by a linear readout grows exponentially with the dimensionality of the information carried by the neurons (Fusi et al., 2016). Increasing the level of mixed selectivity in SUB might therefore be a mechanism by which the hippocampal formation makes relevant output more accessible to downstream target regions.

With these advantages of mixed selectivity in mind, we asked if SUB modifies output from the hippocampus by combining, in individual neurons, multiple features of the navigation experience, in ways that depend on current task goals. Considering that much of the output from CA1 is also passed on to SUB, we performed simultaneous *in vivo* electrophysiological recordings in these regions in rats performing either random foraging or a memory-based spatial navigation task. We compared representations in SUB and CA1 for three navigational covariates: position, head direction and speed. We report that individual neurons in SUB combine these behavioral covariates more extensively than their counterparts in CA1, and more strongly during goal-directed spatial navigation than during free foraging. This coding scheme allowed each of the navigational variables to be decoded more accurately from SUB than CA1, providing regions downstream of SUB with broad spectra of information even from limited numbers of SUB output cells.

## Results

### Anatomical location of recording electrodes

In order to understand what the SUB adds to the navigational output of the hippocampus, we performed extracellular recordings in CA1 and SUB (Figure 1A). All tetrodes were placed in the dorsal one-third of each region (see sectioning plane in Figure 1A). In Nissl-stained coronal sections, CA1 was identified as the narrow, densely packed layer of small pyramidal cells that extends from CA2 (with a less compact cell layer and larger neurons) to SUB (defined as the thicker, more diffuse layer of medium-sized neurons located dorsomedially to CA1). Moving away from the septal pole, SUB gradually widens, extending all the way medially until an additional granule layer is added ventrally, almost at the midsagittal side of the hemisphere. This additional layer belongs to retrosplenial cortex (RSC; Area 29) in the septal part of the hippocampal formation (RSC in Figure S1) and to the dorsal presubiculum in more temporal parts of the structure (preSUB in Figure 1B and Figure S1A). At the septal pole, CA1 continues medially into the fasciola cinereum (FC in Figure 1B and Figure S1) (Boccara et al., 2015). Recordings from FC were not included in our study. Recordings from nearby SUB were included if the tetrodes were more than 50 um away from FC (SUB* in Figure 1B and Figure S1; see ‘Histology and reconstruction of tetrode placement’ in STAR Methods).

**Figure 1.**
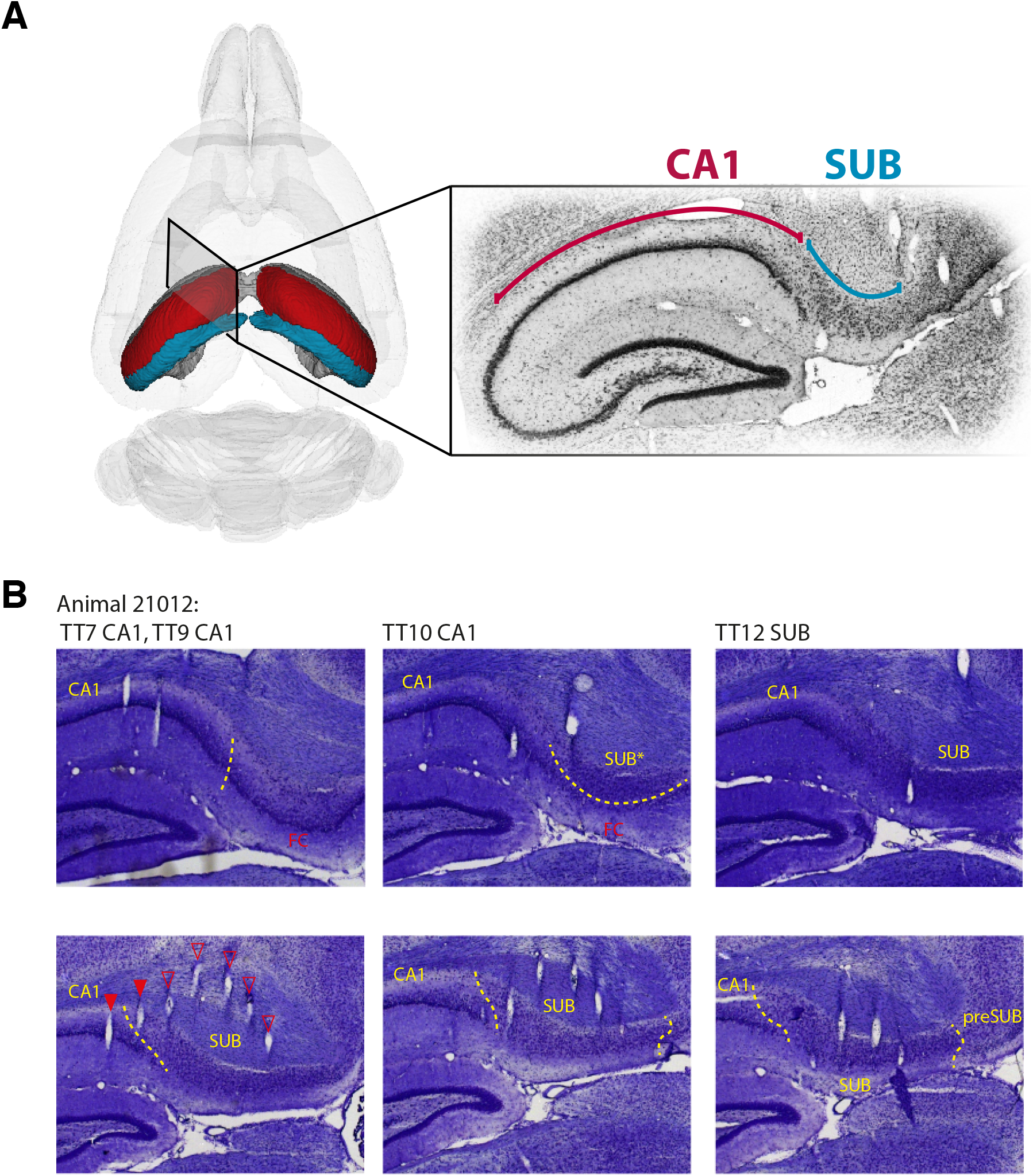
Recording locations in CA1 and SUB. (A) Left side: Schematic of the hippocampal formation, with the CA1 in red and the SUB in blue. Right side: Nissl-stained section showing the arrangement of CA1, with its dense layer of cell bodies (underneath the dark red line following the curvature of CA1), and the SUB, where neurons are more loosely packed (underneath the blue curved line). (B) Reconstruction of the tetrode locations in a representative animal (no. 21012). A micro-drive with two parallel rows each of 7 tetrodes (TT) (6 recording TTs and 1 reference TT) was implanted along the transverse axis of the hippocampus. For unambiguous reconstruction of tetrode locations, the brain was sectioned at a 45-degree angle between the sagittal and coronal planes, parallel to the rows of tetrodes. Filled red arrows indicate tetrode traces near the estimated recording location (usually at the end of the tetrode track), while empty red arrowheads indicate traces of tetrode tracks that are visible in this section but are above (or below) the locations on the track where the recordings were conducted. The TT number above each Nissl section indicates the identity of tetrodes with filled arrowheads, arranged from left to right on the section. Dashed yellow lines indicate borders between SUB and neighboring regions like presubiculum (preSUB) and faciola cinerata (FC). For recording locations in SUB that were dorsal to the FC (labeled with SUB*) or the border between CA1 and SUB, special care was given to include data only from neurons that could confidently be assigned to be 50 μm or more away from the border.

We isolated 760 putative principal neurons from recordings in 8 rats (see STAR Methods). Only cells with clear cluster separation from other background and other neurons were accepted (Figure S2A). Among the accepted neurons, 421 were located in SUB and 325 in CA1 (Table S1B). In SUB, the recording electrodes were distributed quite evenly along the proximo-distal axis (from CA1 to FC, RSC or preSUB) while in CA1 there was a bias for the electrodes to be positioned in mid to distal parts of the subfield, i.e. nearer the SUB boundary (only 2 out of 18 tetrodes were located near the proximal end of CA1, near CA2 (tetrodes 1 and 11 in animal 24101 in Figure S1). However, the number of neurons recorded was distributed more evenly along the proximo-distal axis (approximate neuron numbers in the respective regions: 35 in proximal CA1, 223 in mid CA1, 49 in distal CA1, 292 in proximal SUB and 136 in distal SUB; arbitrary boundaries dividing subfields in three equal bands).

### Position coding in CA1 and SUB

Recordings were undertaken while rats navigated freely in a dimly-lit 1.5 m square environment polarized by a cue card on one of the walls. All distal cues were kept constant (including the cue card) such that after pretraining, when the recording started, the rats were highly familiar with every location in the environment. In agreement with previous studies (Kim et al., 2012; Sharp and Green, 1994), the average firing rate of principal neurons was significantly higher in SUB (4.44 ±0.14Hz) than in CA1 (1.7 ±0.12 Hz, p = 4.6e-53, Welch’s test; Figure 2A and S2A, left panel). Similarly, as in previous studies (Sharp, 1997, 2006; Sharp and Green, 1994), this difference manifested in the shape of spatial firing fields in the two regions. While CA1 cells had sharply defined firing fields (for examples see Figure 2A, left column; 95 % of spikes in CA1 neurons fell into 25.7 ± 1.1% of the spatial bins), SUB neurons tended to fire continuously across the environment, with less distinguishable firing fields (Figure 2A, right column); 95% of the neurons’ spikes fell into 56.3 ± 0.7% of the spatial bins). In line with other findings (Lever et al., 2009; Stewart et al., 2014), a large proportion of the position-modulated neurons in SUB showed boundary-vector-like activity, meaning that their firing fields were elongated parallel to a wall of the environment (see Figure 2A and Figure S2B). The information rate of the neurons, or their amount of position information per time interval (Skaggs et al., 1993), was similar in CA1 and SUB (Figures 2C, top, and S2D; medians and median absolute deviation (MAD) are 0.47 ± 0.27 bits/s in CA1 and 0.42 ± 0.26 bits/s in SUB), whereas information content, or the specificity of a neuron’s individual spikes for position (Skaggs et al., 1993), was markedly smaller in SUB than in CA1 (Figures 2C, bottom, and S2E; median and MAD for CA1: 0.84 ± 0.49 bits/spike, for SUB: 0.11 ± 0.06 bits/spike).

**Figure 2.**
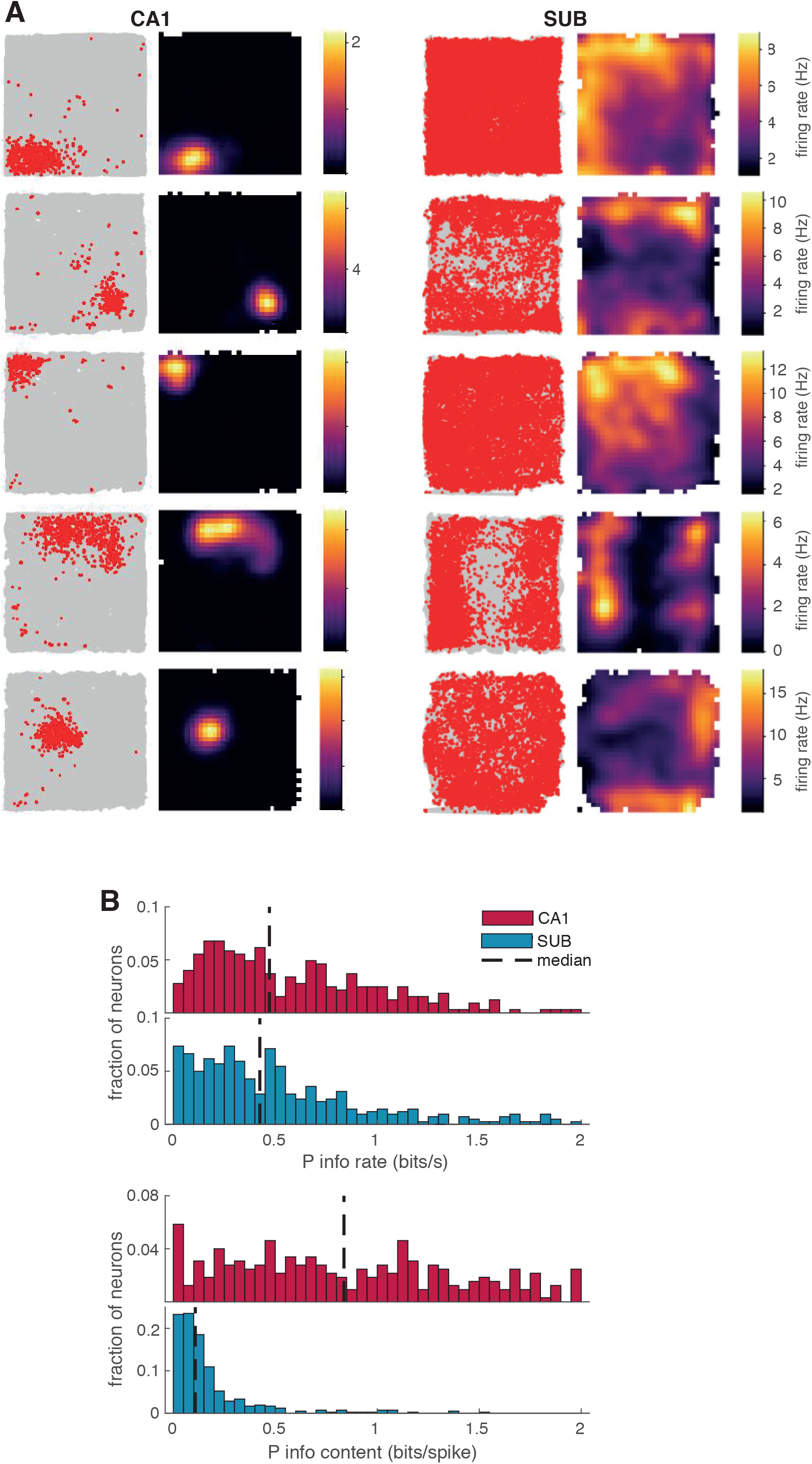
Difference in position coding in CA1 and SUB. (A) Example path plots (left) and rate maps (right) from neurons in the CA1 (left column) and SUB. Path plots show the path of the animal during the entire session in gray and the emitted spikes of the respective neuron overlaid in red. Color bars to the right of each rate map indicate firing rate. (B) Frequency distributions showing position information rate (top) and information content (bottom) for all neurons recorded in CA1 (red) and SUB (blue) in all animals. The dashed line indicates the median of each distribution.

The difference in information content but not rate raises the question of what benefits are conferred by sacrificing information per spike during the processing step from CA1 to SUB. Potentially, SUB may integrate positional information over longer time scales than CA1, which would allow individual SUB neurons to combine input from CA1 with information from other sources (including inputs encoding variables besides position). Therefore, we investigated whether information about other behavioral variables is expressed in spike trains of SUB neurons.

### Representation of multiple covariates including head direction and speed in SUB and CA1

Previous studies found that subsets of CA1 and SUB neurons show some degree of modulation by navigational variables like head direction modulation and running speed. In CA1, place cells may respond to head direction specificity inside their place fields (Acharya et al., 2016; Langston et al., 2010; Leutgeb et al., 2000). Weak head direction tuning has also been reported for spatially modulated neurons in SUB (Sharp and Green, 1994). Similarly, place cells in CA1 respond to some degree to running speed whereas speed tuning has not been reported in SUB to our knowledge (Czurkó et al., 1999; Kropff et al., 2015; McNaughton et al., 1983).

With these observations in mind, we set out to compare quantitatively, in the same recordings, the extent to which cells in CA1 and SUB represent navigational covariates besides position, like head direction and speed. The head-direction tuning curves of the recorded neurons revealed varying degrees of modulation by head direction, in both CA1 (Figure 3A, 2^nd^ column) and SUB (Figure 3B, 2^nd^ column). Consistent with previous reports in CA1 (McNaughton et al., 1983; Skaggs et al., 1993), the information rate in CA1 neurons was much lower for head direction than for position (Figure 3C, median and MAD of the CA1 population was 0.04 ± 0.04 bits/s). In contrast, we found that SUB neurons express comparable information rates for both head direction and position (median and MAD for head direction 0.22 ± 0.15 bits/s). CA1 and SUB neurons showed similar levels of information content (medians and MAD for information content of 0.05 ± 0.05 bits/spike for CA1 and 0.07 ± 0.06 bits/spike for SUB).

**Figure 3.**
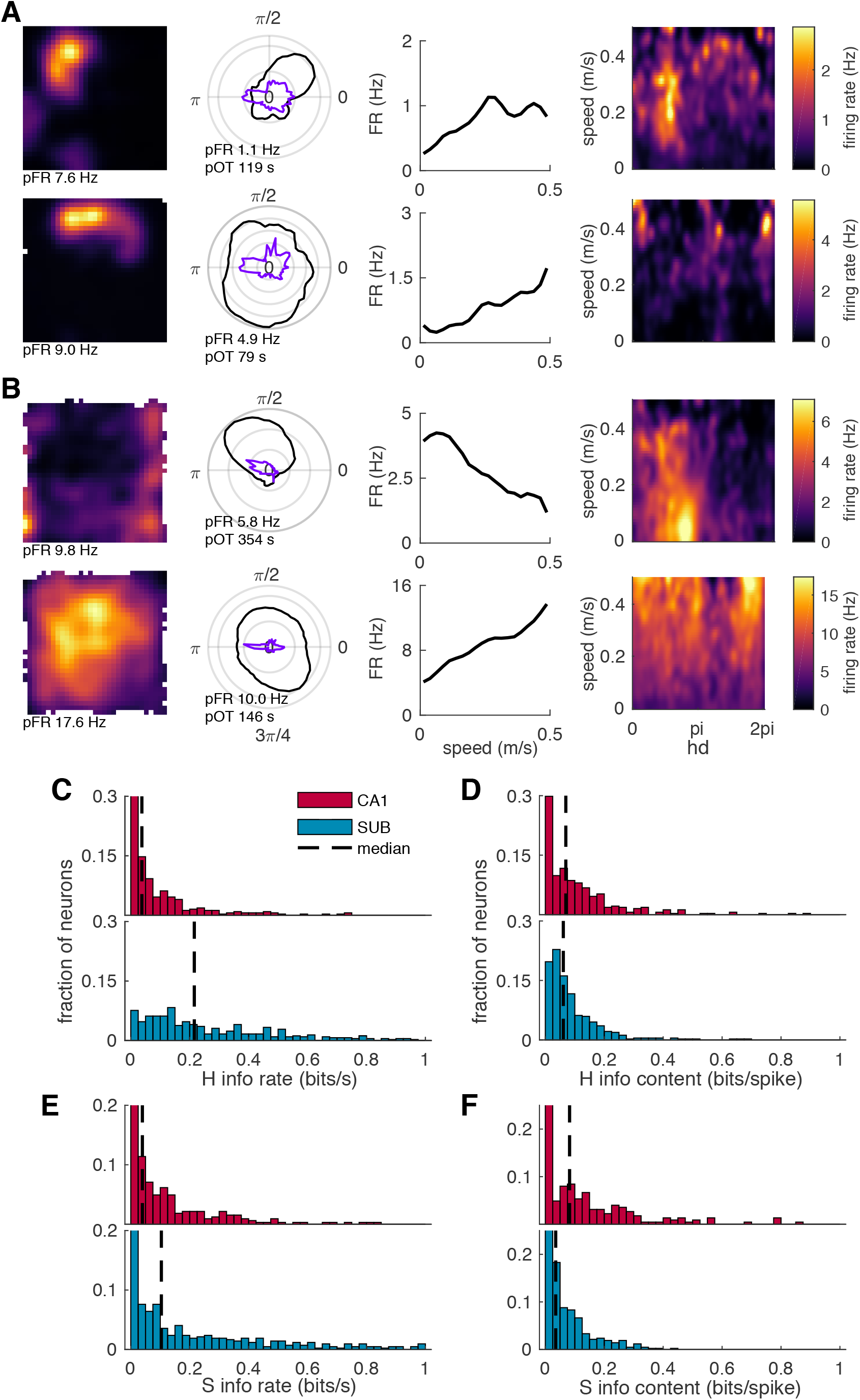
Information rate for head direction and speed is higher in SUB than CA1. (A-B) Example neurons from CA1 (A) and SUB (B). Representative firing rate maps (first column; as in Fig. 2), head direction tuning curves (second column; polar plots showing firing rate FR as a function of head direction), speed tuning curves (third column; linear plots showing firing rate as a function of speed), and head direction versus speed rate maps (fourth column; firing rate color-coded as a function of head direction and speed). Peak firing rates (pFR) are indicated for rate maps and head direction tuning curves. In the second column, the black curve shows firing rate whereas purple shows occupancy time. Peak occupancy time (pOT) is indicated. (C-D) Frequency distribution showing scores for head direction (H) information content (in C) and information rate (in D) across all neurons recorded in CA1 (red) and SUB (blue). Dashed lines indicate the median of each distribution. (E-F) Distribution of speed (S) information content (in E) and information rate (in F) across all neurons recorded in CA1 (red) and SUB (blue). Dashed lines indicate the median of the distribution.

A similar pattern could be seen for speed information rates and contents (Figure 3E and 3F). CA1 and SUB neurons had similar information content for speed, while information rates were significantly higher in SUB than CA1 neurons. The median and MAD for speed information rate were 0.04 ± 0.04 bits/s in CA1 and 0.10 ± 0.10 bits/s in SUB (Figure 3E). Median and MAD for speed information content were 0.07 ± 0.07 bits/spike in CA1 and 0.04 ± 0.04 bits/spike in SUB (Figure 3F). Thus, it appears that SUB neurons exhibit a higher information rate relative to information content for all three covariates (position, head direction, and speed), while CA1 cells show higher information content relative to information rate for head direction and speed.

### Modeling spike rate with a generalized linear model reveals mixed selectivity in SUB

Measuring information rate separately for each different covariate (position, head direction and speed) entails some important limitations. These three behavioral covariates are strongly interdependent, meaning that an effect which is selectively tuned to one covariate might manifest spuriously when measured in relation to a different covariate (Acharya et al., 2016). This undesirable effect may result in both false-positive and false-negative tuning outcomes. One solution to this problem is to employ multiple-variable statistical methods, such as the generalized linear model (GLM). The GLM considers multiple covariates simultaneously, and thus allows the influences on a cell’s activity to be ascribed in a principled manner to the covariates that provide the strongest prediction.

Therefore, we extended the analysis by using a Poisson GLM framework (McCullagh., 1989, Hardcastle et al., 2017; see Methods section) to investigate tuning of the neurons to position (P), head direction (H) and speed (S), while accounting for any deceptive correlations between the covariates induced by sampling. Alongside the behavioral covariates (P, H, S), we also included two covariates representing basic influences on neural activity: ensemble activity (E), defined as the z-scored spike count summed over all other neurons on the same tetrode as the cell in question, and the theta phase (T) of the filtered local field potential (see Methods section). These two covariates were treated as ‘internal covariates’ which helped fitting the model to the spike counts but did not account for the modulation of SUB neurons by the animals’ behavior. While they were included in the models throughout the study to optimize fitting of every neurons’ spike count, we focused our attention on the behavioral covariates P, H and S (for extended discussion see Methods section). All the covariates (P; H; S; E; T) were binned along their respective dimension (30×30 bins for P, and 10 bins for each H, S, E and T), while continuity in the predicted tuning curves was enforced via a smoothness prior (see Supplementary Methods). All models resulting from specific subsets of the covariates (ranging from single-covariate models, e.g. only P, to the most complex model containing all five covariates, PHSET) were trained and tested via 10-fold cross-validation for every cell. Bin size and smoothness prior were treated as hyperparameters and selected by optimizing the cross-validated log-likelihood for each single covariate model (Figure S3). For each neuron, the full model GLM (PHSET) yielded tuning curves for P, H, and S (Figure 4A – C, panels in the lower row labeled with MODEL), which qualitatively reproduced the tuning curve for the data (Figure 4A – C, panels in the upper row labeled with DATA).

**Figure 4.**
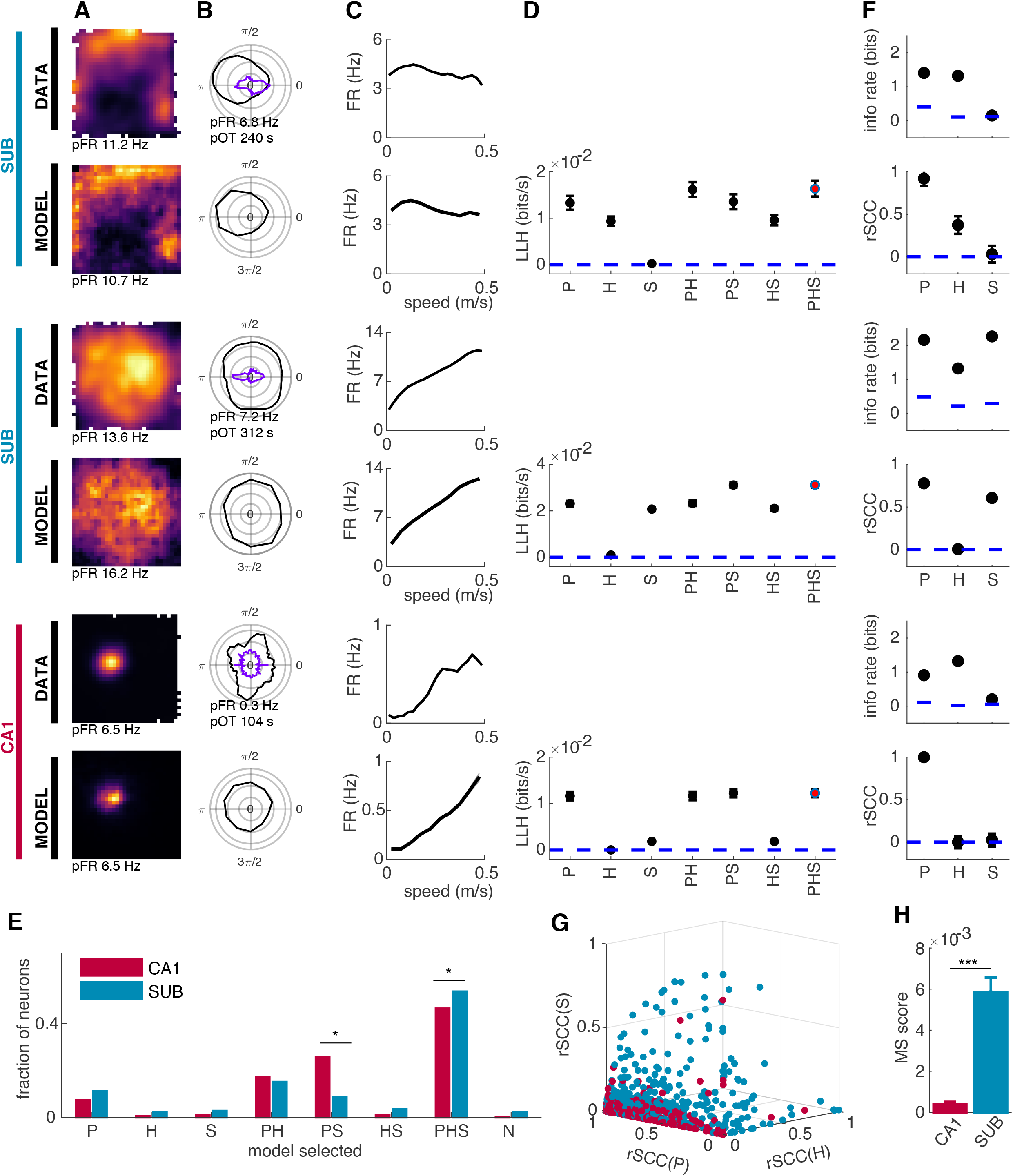
Modeling the data with a generalized linear model (GLM) shows contribution of position, head direction and speed to firing rates of individual neurons. (A-C) Spatial rate maps and tuning curves for head direction and speed for three example neurons generated from data (upper panels labeled with DATA) or from spike trains reproduced with the GLM (lower panels labeled with MODEL). Rate maps, polar plots and speed tuning curves as in Figure 3. Note minimal tuning to head direction in the CA1 neuron (peak firing rate ~0.3 Hz). (D) Log-likelihoods (black circles) resulting from fitting the spike train of each neuron in A-C with models encoding one covariate - position, head direction or speed (either P, H or S) - or with combined models containing two or more covariates. Models that performed significantly better than all less-complex models are indicated in red. Error bars are S.E.M. across ten cross-validation folds. Log-likelihoods from models containing ensemble activity (E) or theta (T) as additional covariates (Figure S4) were collapsed with simpler models containing P, H and/or S (for example PE is collapsed with P). (E) Frequency distribution showing fractions of neurons from CA1 (red bars) and SUB (blue bars) selecting the different models. With the exception of the PHS and the PS models, models were not selected at significantly different frequencies by the two populations. PS, however, was selected by 88.9 % CA1 compared to 81.3 % of SUB neurons and PHS was selected by 40.4 % of the neurons in CA1 as compared to 53.7 % in SUB (difference > 99.9 percentile of a spike time-shuffled distribution for both PS and PHS). (F) Information rate (info rate; upper panels) and relative single covariate contribution (rSCC; lower panels) for the neurons shown to the left in A-D. Note that even if all three neurons in D select the most complex model (PHS), the rSCCs indicate that the firing of the CA1 neuron is predominantly determined by the animal’s position while the firing of the two SUB neurons is determined by P and H or P and S, respectively. (G) Three-dimensional plot of the rSCCs of P, H and S. Each neuron is represented by one datapoint in red for CA1 and in blue for SUB. Note abundance of SUB neurons with high values for more than one rSCC. (H) Mixed selectivity score (MS score) defined as the product between rSCC(P), rSCC(H) and rSCC(S).

Model selection was performed in a forward stepwise fashion, starting from single covariate models and adding one covariate at a time using a non-parametric test and a 5% significance level (for details see Methods). Except in Figure S4H, models containing internal covariates E and/or T where pooled with the similar models without this covariate (e.g. neurons best modeled by the PHT, PHE and PHTE were counted to the group of the PH model). In this way we determined for every neuron which model provided the best fit to the neuron’s firing properties in the data (red dot in Figure 4D), which in turn allowed us to assess model performance at the population level (Figure 4E). The proportion of cells for which the most complex model (PHS) performed significantly better than any simpler model was higher for SUB than CA1 (53.7 % in SUB vs. 46.5 % in CA1; Figure 4E, difference > 99.9 percentile of a distribution of shuffled data where cells were randomly assigned to both anatomical regions). Conversely, the proportion of neurons best fitted by the PS model was significantly lower in SUB than in CA1 (25.9 % in CA1 and 8.8 % in SUB, difference > 99.9 percentile of a shuffled distribution; Figure 4E). These findings may suggest that both CA1 and SUB cells express high levels of mixed selectivity but the degree of mixing is stronger in SUB.

However, one caveat of the model-selection analysis reported above is that it only allows us to conclude that a number of CA1 and SUB neurons were modulated by position, head direction and speed in a statistically significant manner, while it did not inform us on how strong this modulation was. As a next step we therefore investigated how strongly each neuron’s firing rate was determined by either an individual covariate (position, head direction and speed) or a combination of them. Using the GLM framework, we quantified the relative contribution of every single covariate (rSCC) by taking the difference between the log-likelihood (LLH) of the selected model and the LLH of the model with the respective covariate removed. This term was then divided by the square root of the sum of the squares of all differences. The contributions of covariates not used in the selected model were set to zero (see Equation in methods section ‘relative single covariate contributions’). rSCC was in many cases correlated with information rate for the respective covariate (Figure 4F; information rate in top row and rSCC in bottom row for each neuron) but occasionally there were also large differences (see covariate H in the third example in Figure 4F). While the measure of information rate is biased due to correlations between covariates, rSCC is derived from the GLM framework, allowing us to simultaneously quantify the contribution of multiple covariates. This leads to a more differentiated estimate of the contribution of individual covariates to the cell’s firing rate (Figure 4F, see rSCC versus information rate in the two SUB neurons).

At the population level, plotting the rSCC for CA1 and SUB neurons separately (Figure 4G red and blue circles, respectively) revealed that CA1 neurons were largely distributed along the axis for position, showing very little contribution of head direction and speed to their firing pattern, despite the fairly large number of cells that selected the PHS model in the above analyses. In contrast, SUB neurons were scattered throughout the entire space, indicating the differential combinations of the three covariates determining the firing of the neurons. This suggested that, albeit showing a high level of mixed selectivity in terms of model selection, CA1 neurons singled out one covariate - position - that dominated their tuning properties, whereas SUB neurons more strongly combined multiple covariates, expressing a conjunctive code. In order to quantify this coding difference between CA1 and SUB, we introduced a mixed selectivity score (MS score). The MS score is the product of all rSCC’s in each neuron (see methods section: ‘mixed selectivity score’). This results in a maximal score for neurons that have equal contributions of position, head direction and speed to their firing rate. The average MS score for SUB neurons was 5.8e^−3^ ± 7.1e^−4^, one order of magnitude larger than for the CA1 neurons, which had an MS score of 3.8e^−4^ ± 1.2e^−4^ (Figure 4H; Wilcoxon signed-rank test, p = 6.86e-9). Thus, in contrast to CA1 cells, SUB cells integrate information regarding position, head direction and speed in a balanced fashion, resulting in a highly mixed selective population in which every neuron expresses differential contributions of the three covariates.

### Mixed selectivity in SUB is task-modulated

Spatially modulated cells are often recorded while animals forage for randomly spread cookie crumbs in an open field, as in Figures 2-4. To explore the function of mixed selectivity in the SUB further, we probed it in a behavioral situation where the animal had to retrieve its current position, head direction and speed regularly in order to correctly respond to task requirements. We adapted a spatial memory task (ST) from Pfeiffer and Foster (2013) in which the rats alternate between free search and goal-directed navigation. The animals performed this task in the same arena as used for random foraging (open field, OF). Between ST and OF the floor of the arena was exchanged from a black even mat without incisions (OF) to a black mat with a square grid of 1 cm-diameter holes. The walls of the arena and their location in the room with respect to all distal landmarks remained constant (Figure 5A). During the ST, chocolate oat milk was provided alternatingly at a fixed ‘home well’ and in a randomly selected well (‘random well’) of the arena (Pfeiffer and Foster, 2013). The animals memorized the home well location and learned to navigate back to it straight after consuming the randomly placed reward (Figure 5A), as reflected in their behavioral latencies (Figure S4A). There was a marked change in behavior between the two tasks: In the OF, the animals roamed unpredictably from one place to another in search for randomly spread crumbs, while in the ST, they either attentively checked one well after the other to find the reward in the random location (targeted search phase), or they actively navigated back to the previously memorized home well (memory phase).

**Figure 5.**
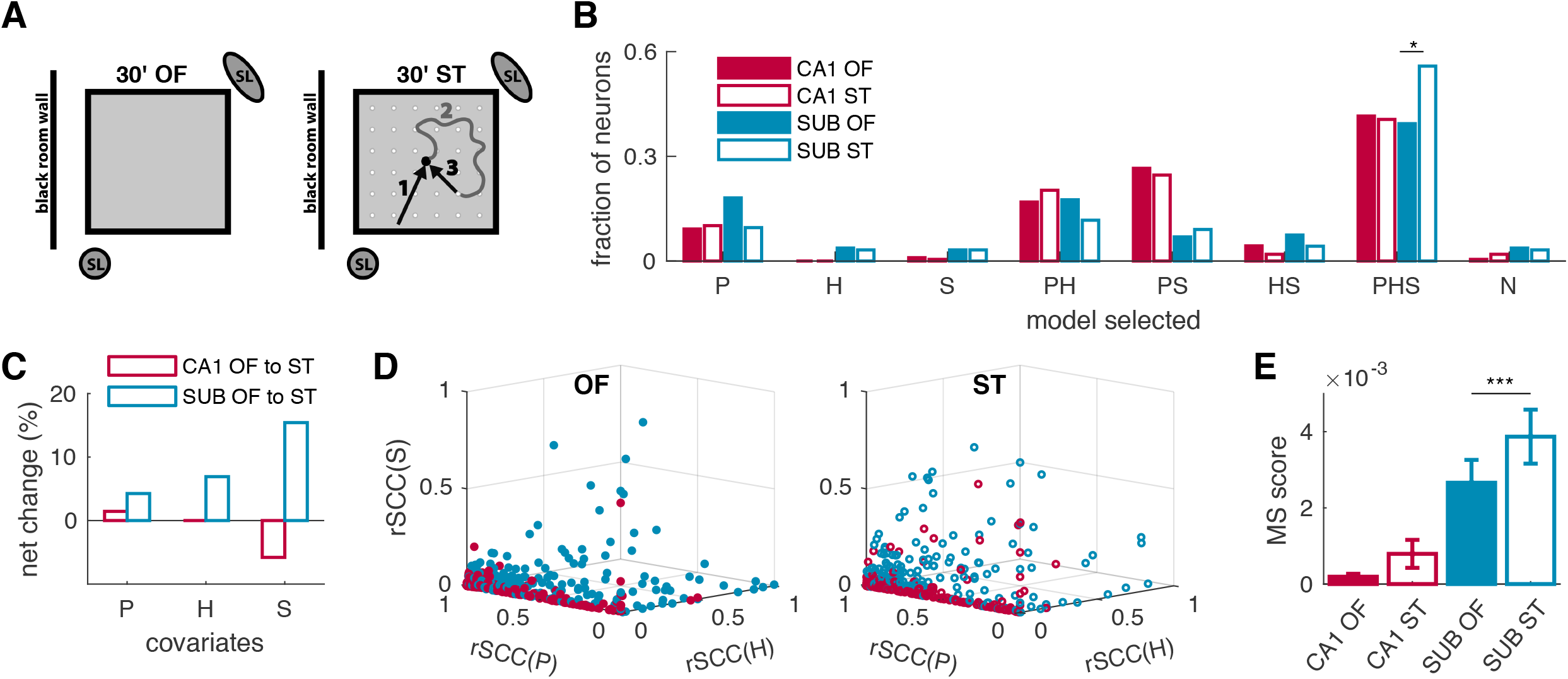
SUB neurons exhibit more mixed selectivity in the spatial task (ST) than during random foraging in the open field (OF). (A) Schematic representation of the OF and the ST. Spatial landmarks (SL) and location of the enclosure with respect to room and experimenter are constant between the two tasks. (B) Model selection in CA1 (red bars) and SUB (blue bars) for recordings in the OF (filled bars) and the ST (empty bars). The model selection process is significantly different in the two instances in SUB: In SUB but not CA1, the population of neurons best fitted by the PHS model is significantly increased, in response to the task change from OF to ST. (C) Percentage of neurons that add or remove the covariates P, H and S after the task change. (D) rSCC for position, head direction and speed for each neuron in the OF (left graph) and the ST (right graph) for the CA1 (red datapoints) and the SUB (blue datapoints). (E) MS score in CA1 (red bars) and SUB (blue bars) in the OF (full bars) and the ST (empty bars).

In order to understand whether neuronal tuning to the selected behavioral covariates adapts to task context, we quantified position, head-direction and speed parameters in the two tasks. Even though the floor was exchanged between tasks, we only observed small effects on the location of the firing in individual CA1 and SUB neurons (examples in Figure S4B). Albeit this, the correlation between tuning curves for position, head direction and speed where lower across task conditions than within task (Figure S4C). However, the levels of information rate and information content stayed well within the range of the respective region (i.e. the CA1 population showed always at higher levels of information content compared to SUB while SUB showed larger information rate throughout; see Figure S4D and E).

Using the previously introduced generalized linear model, we next set out to quantify the level of mixed selectivity in the network across the two task situations. For CA1, the distribution of selected models was similar between OF and ST (Figure 5B, red bars, no changes in model selection were higher than the 99.9^th^ percentile of a shuffled distribution). In contrast, the proportion of SUB neurons for which the most complex model (PHS) performed significantly better than the others, increased from OF to ST (Figure 5B). While other changes in model selection between OF and ST were not significant, the increase from 39.4% to 55.9% of neurons best fitted with the PHS model was significantly beyond the 99.9^th^ percentile of a distribution of shuffled data (Figure 5B). A total of 2.3% and 1.4 % of the neurons in SUB and CA1, respectively, were tuned to position only during ST and not in OF. In SUB 6.9% and 15.4% more principal neurons were tuned to head direction and speed, respectively, in ST but not in OF. In CA1 5.8 % of the neurons lost their tuning for speed after transitions from OF to ST (Figure 5C). When considering tuning to ensemble activity and theta (which were always fitted in parallel to the other three covariates), it was apparent that the activity in the largest proportion of SUB neurons (35.6 % in OF and 51.6% in ST) was best fitted by all 5 covariates (Figure S4F) while in CA1 the largest proportions of neurons (42.5 % in OF and 41.5 % in ST) were best fitted by a 4 covariates model (Figure S4F). The latter largely consisted of the PHET and the PSET models (see Figure S4H). The task change did not affect the overall likelihood of the best-performing model (Figure S4G), indicating that the fitting procedure performed similarly well in both behavioral situations.

The increase in mixed selectivity in SUB was reflected in the relative contribution of each single covariate to the firing rate of the neurons. When plotting the rSCC separately for OF and ST, it appeared that during the ST condition the SUB datapoints cluster more towards the center of the 3D plot, indicating a more equal contribution of the three covariates to the firing pattern of the cells (Figure 5D). To quantify this phenomenon, we calculated the mixed selectivity (MS) score introduced before. Taking the product between all rSCC s of the three covariates showed that, after transitions from OF to ST, there was a significant increase in mixed selectivity in the SUB (from 1.3e^−3^ ±0.58e^−3^ in OF to 2.8e^−3^ ±0.42e^−3^ in ST, p = 0.00018, Wilcoxon signed-rank test). In CA1, the mixed selectivity score was low in both task conditions (0.02e^−3^ ±0.02e-3 in OF to 0.07e^−3^ ±0.01e-3 in ST, p = 0.47, Wilcoxon signed-rank test, Figure 5E). In summary the task change led to a clear increase in mixed selectivity in SUB, suggesting that, in contrast to CA1, the SUB code flexibly adapts to altered behavioral situations, changing its coding regime according to task requirements.

### High decoding accuracy for position, head direction and speed in SUB

The previous analyses using information-theoretic measures and GLM-based model selection established that the responses of neurons in CA1 and SUB carry information about multiple covariates, and that a statistical model requires the mix of these covariates to best explain the data, more so in SUB than CA1 and more influenced by the task in SUB than in CA1. However, it is not straightforward to directly translate these results into what can actually be read out from these codes in upstream cell populations. In order to understand what the large amount of mixed selectivity in SUB might add to the hippocampal output, we therefore next tried to decode position, head direction and speed simultaneously from cell populations in either CA1 or SUB.

Methodological constrains limited the number of simultaneously recorded neurons in subiculum (the recording range was 3 to 5 neurons per SUB tetrode). In order to study decoding performance as a function of population size and extrapolate the decoding power of the two regions across tasks for a large number of neurons, we exploited the GLM framework to resample all the recorded neurons over two concatenated session for ST and OF respectively (see methods section ‘*resampling*’). This allowed decoding of position, head direction and speed at the same time. Decoding of all three covariates was more accurate when using data from SUB than CA1 (Figure 6A and B). To assess the significance of this difference we shuffled the neural populations with respect to their anatomical origin and computed the differences between two sets of randomized selections of neurons (ensuring equal probability to select neurons from the CA1 or the SUB cell population; Figure S7). This shuffling analysis showed that the difference in decoding accuracy deviated significantly from a distribution of shuffled differences (p < 10e^−3^) for decoding position, head direction and speed in the open field and the spatial task. While smaller decoding errors for SUB compared to CA1 were found across all population sizes, these differences were only significantly different from the differences in a shuffled distribution for population sizes larger than 100 neurons for position, larger than 20 neurons for head direction, and larger than 50 neurons for speed (for population sizes of 5, 20, 50, 100 and 150, respectively, significance values were, for position: p = 0.66, 0.61, 0.32, 0.005 and 0; for head direction: p = 0.02, 0, 0, 0 and 0 and for speed: p = 0.595, 0.240, 0.042, 0.033, 0.009).

**Figure 6.**
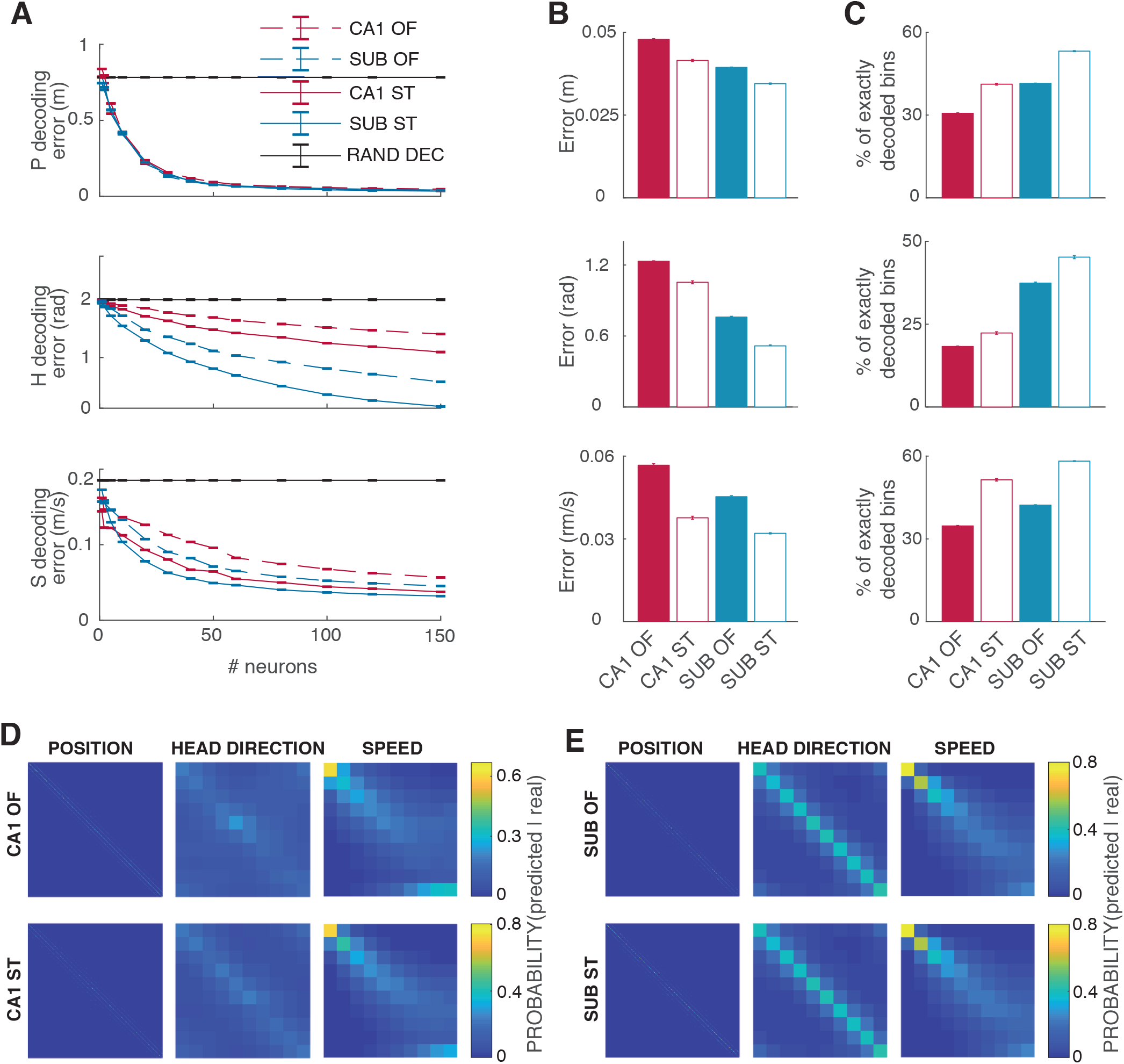
More accurate decoding of position, head direction and speed from a fixed number of cells in SUB compared to CA1. (A) Decoding of position (top panel), head direction (middle panel) and speed (bottom panel) for different numbers of resampled neurons. For all three covariates, decoding errors are smaller in SUB (blue) than CA1 (red) and smaller in the ST (full lines) than in the OF (dashed lines). (B-C) Decoding error (B) and percentage of perfectly decoded bins (the number of time bins in which the decoded was the exact same as where the animal roamed at this instance; C) in a population of 150 neurons, shown separately for position (top panels), head direction (middle panels) and speed (bottom panels). Note smaller decoding error for position in SUB than CA1. Error bars correspond to one S.E.M. over 20 samples of the identities of the resampled cells. (D-E) Confusion matrices for decoding in CA1 (D) and SUB (E). Columns show decoded covariates: position (left), speed (middle) and head direction (right); rows show tasks: OF (top) and ST (bottom). Within each confusion matrix C, rows identify decoded covariate bins, while columns identify bins for the covariate in the data, such that position confusion matrices are 900×900 and head direction or speed matrices are 10×10. The color-coded element C_ij_ of the confusion matrix represents the fraction of time bins where the decoded covariate bin is i given that the actual covariate bin was *j*. Bright yellow colors indicate maximum overlap while dark blue colors indicate no overlap between decoded and actual covariate.

To better understand the difference in coding in SUB versus CA1, we also estimated the relative amount of perfectly decoded bins (PDB). The PDB is the number of time bins in the time series for which the decoder settled on the exact bin of position, head direction or speed where the animal was in during that time bin. For all three covariates, position head direction and speed, the amount of PDB was higher when using SUB as opposed to CA1 neurons (Figure 6C; these differences were significantly different from randomized distributions). This indicates that, for all three covariates, the neural code in SUB allowed more accurate decoding than the one from CA1, suggesting that the mixed selective rate code of the SUB is an effective way of transmitting signals to downstream target regions.

Comparing decoding accuracies across the two tasks showed that the decoding error for position, head direction and speed were significantly reduced in the ST versus OF (Figure 6A and B, p < 10e^−3^ compared to a shuffled distribution). Decoding accuracy was higher in the ST than the OF also when counting the number of PDBs (Figure 6C). While this was true for both CA1 and SUB, overall the SUB reached a significantly lower number of decoding errors as well as higher proportions of PDB for position head direction and speed (Figure 6A and B, p<10e^−3^ in shuffling test of significance). In order to ensure that the improved decoding in the ST did not result from a behavioral bias, we plotted confusion matrices displaying decoding accuracies for the different bin identities (Figure 6E and F). For head direction and position, the confusion matrices did not show any bias towards disproportionally better decoding in certain bin clusters of the data than others. For speed decoding, however, the confusion matrix showed a bias for better decoding in the lowest speed bins (0-10 m/sec; Figure 6E and F, rightmost column). This bias was present for decoding from CA1 and SUB (Figure 6E and F top and bottom row, respectively) and for the OF and ST and might be a result of the disproportionally high sampling in these bins. Furthermore, even though the average speed was similar between OF and ST, the distribution was skewed towards lower and higher values (meaning that in the ST the animal would be more often entirely immobile in order to consume the reward and then run faster to the next well). We therefore tested if decoding was affected by these differential speed distributions. By sampling the speed equally for decoding from OF and ST, we could show that decoding of all three covariates – position, head direction and speed – was still improved in the ST over the OF (Figure S5A-C). This led us to conclude that improved decoding accuracy did not result from of spatial sampling bias but from other functional differences, attributable to the spiking pattern forming the output of CA1 and SUB during the ST.

One might argue that the superiority of decoding in the SUB data might be affected by our choice of GLM as a modeling framework as the goodness-of-fit of a model always depends on the choice of the measure used. However, in our data, neural activity in CA1 and SUB are equally well modelled by the GLM when the goodness-of-fit is measured by the difference in the likelihood of the best model and the average firing rate model (Figure S5E). Furthermore, when the goodness-of-fit is measured with explained deviance, the model fit is even better in CA1 than in SUB neurons (Figure S5F). Additionally, the task choice does not appear to affect the model performance. We therefore consider it unlikely that the decoding results should be merely a consequence of our choice of the GLM as a modelling framework. Instead the higher decoding accuracy of all navigational covariates points to an important role of SUB of integrating multiple information streams from the hippocampal formation and providing an accurate navigational code to downstream regions.

## Discussion

Downstream regions of the hippocampal formation rely on output from the CA1 and SUB subfields during behaviors when navigational and mnemonic information is relevant. While spatial coding in CA1 has been investigated extensively in place cells, it has remained elusive how SUB transforms the positional code. Here, we record neural activity during two different spatial behaviors and show that SUB provides a mixed selective code from which three correlates of navigation - position, head direction and speed - can each be decoded at higher accuracies, from a similar number of neurons, than from the CA1 output. The presence of mixed selectivity is consistent with an early report of place and head direction tuning in the same SUB cells (Sharp and Green, 1994) but takes the finding further by quantifying the mixing of three navigational variables and by showing that decodability is enhanced compared to CA1. Our experiments further demonstrate that mixed selectivity in SUB increases with navigational task demands, suggesting a behaviorally dependent information transfer to downstream regions. The activity of individual SUB neurons, which are informative on a broader spatial scale (and consequently, a longer temporal scale) than CA1 cells, may represent an integration of inputs from CA1 with information from other areas. In the present work, the combination of covariates into a mixed representation is demonstrated for three navigational parameters: position, head direction and speed. However, it has been shown that hippocampal neurons respond also to other covariates like odors (Dusek and Eichenbaum, 1997), texture (Wood et al., 2000) and time (Eichenbaum, 2017; Fortin et al., 2002; Hampson and Deadwyler, 2003; Kesner et al., 2002). This raises the possibility that SUB neurons combine a wide range of covariates, resulting potentially in a behavior-dependent mixed code for retrieval of a broad spectrum of experiences in downstream regions.

In an information theoretical analysis, Kim et al. (2012) showed that SUB is well suited to transmit information about the animal’s position using fewer neurons than the CA1 region. We show here, with experimental data, that we can indeed decode position at higher accuracy from SUB data than from CA1, when the number of cells is the same. A similar enhancement was seen in SUB for decoding of head direction and speed. For position, decoding can be extremely accurate also in CA1 when a high number of neurons is available (Pfeiffer and Foster, 2013; Wilson and McNaughton, 1993) but it is unclear if the density of connectivity to distant downstream regions is high enough to integrate over such large arrays of inputs. As SUB is the major receiver of CA1 output (Cappaert et al., 2015) projection density to this region might allow SUB to integrate over a large array of neurons in order to translate the sparse code from CA1 into a code that can be decoded accurately from a lower number of neurons at the next stage. Our study suggests that a major function of SUB may be to broadcast the hippocampal signal for more efficient readout in distant brain areas. A recent study by Stringer et al. (2019) made predictions of potential relevance for SUB as a long-range distributor. One notable prediction was that neural responses in an ideal population code should not change sharply in response to small changes in the stimulus value. By this measure, the CA1 code would be a poor candidate for long-range distribution, because its sparsity means that very nearby locations are represented near-orthogonally by the CA1 population. By comparison, SUB, with larger spatial receptive fields than CA1, could provide a population code which varies more smoothly with the animal’s movement and so would make it more efficient as a code for long-range distribution.

Although position, head direction and speed are represented in both CA1 and SUB neurons, combinatorial expression of those variables was a lot more common in SUB neurons. Mixed selectivity neurons have previously been found in a number of cortical and subcortical brain areas (Asaad et al., 1998; Freedman and Assad, 2009; Hardcastle et al., 2017; Meister et al., 2013; Rigotti et al., 2013; Rishel et al., 2013), including the hippocampal areas CA1 and CA3 (Acharya et al., 2016; McKenzie et al., 2014; Wood et al., 2000). Here we provide additional evidence that CA1 neurons encode multiple variables simultaneously, but we show further that representations in this subfield are dominated by position while other navigational covariates have a less predominant influence the firing rates. Neighboring SUB, in contrast, displays a more evenly mixed code in that many neurons are rate-tuned to our three navigational covariates – position, head direction, and speed - to almost equal amounts. We also show that at the population level the mixed selective code generated in SUB is more accurate than the output generated in the CA1. As we find little evidence for highly selective neurons in SUB for any of the three covariates in our information theoretical analyses (Figures 1 and 2), it is unlikely that the high precision decoding in SUB populations comes from a few cells which are highly selective for one variable. The fact that mixed selectivity leads to better combined decoding of the spatial covariates can be intuitively understood in the following way. A randomly selected population of uni-selective cells has a reasonably high chance of including only few neurons selective for a given covariate, thus leading to low precision decoding for that variable. On the other hand, at the same population size, a random sample of mixed-selective neurons will have all variables represented in each cell and therefore the information available for decoding for all variables is more evenly distributed across them.

Our recordings in the spatial task further show that neurons in SUB change their level of mixed selectivity depending on the task the animal is involved in. While it has been observed before that SUB neurons change their strength of tuning to movement direction depending on the animal’s behavior (Olson et al., 2016), we show here that task-dependent changes in tuning strength, in environments with high visual similarity, are not limited to direction (head direction in our case) but also encompass speed and position. Higher cortical areas like the prefrontal and the posterior parietal cortex have previously been shown to adapt their mixed selective neural response to different task demands (Mante et al., 2013; Parthasarathy et al., 2017; Raposo et al., 2014; Stokes et al., 2013). The present findings suggest that, in the hippocampal formation, navigational output is modified by increasing the level of mixed selectivity through a processing step in SUB. This likely serves a mechanism for downstream regions to obtain access to a broad spectrum of hippocampal information when it is behaviorally relevant. However, although mixed-selective coding of position, head direction and speed proved to be effective at predicting variation on these variables, we cannot claim that these variables constitute the only neural code of SUB. The models compare only the covariates that are put into them. As SUB has been shown to be important for learning and memory (Cembrowski et al., 2018a; Morris et al., 1990; Roy et al., 2017), it is likely that its activity relates also to behavioral or cognitive variables that were absent from our analysis of navigational behavior.

Previous theoretical work has shown that mixed-selectivity coding schemes offer certain computational advantages over single-variable selectivity. When multiple covariates are combined in varying quantities in the population of neurons projecting to a downstream target region, a downstream neuron can linearly combine an arbitrary subset of the inputs in order to reconstruct the value of a single covariate (Fusi et al., 2016; Ganguli and Sompolinsky, 2012). This means that, with no need for precise prepatterned connections, all the information can be transmitted, allowing downstream regions to integrate different covariates without having to repattern connections. Keeping in mind that neurons from SUB may consist of genetically compartmentalized subpopulations each projecting to only a few selected target areas (Bienkowski et al., 2018; Cembrowski et al., 2018b; Naber and Witter, 1998; Witter, 2006), the mixed code from SUB may be specifically tailored to ensure that a wide range of covariates computed in the hippocampal formation is accessible to distant projection areas, despite the limited number of SUB cells that may project there. While the widespread projections from CA1 provide position information to a large array of downstream areas, the SUB output ensures the inclusion of a broader spectrum of task-relevant information in the hippocampal output, encoded in ways that can be integrated efficiently by downstream target regions.

## Methods

### Subjects

Data were collected from eight male Long Evans rats, which were experimentally naïve and 3–5 months old (350–600g) at the time of implantation. The rats were group housed with 3–8 of their male littermates prior to surgery and were singly housed in large Plexiglas cages (45×44×30cm) thereafter. The rats were kept on a 12h light/12h dark schedule, and humidity and temperature were strictly controlled. The experiments were performed in accordance with the Norwegian Animal Welfare Act and the European Convention for the Protection of Vertebrate Animals used for Experimental and Other Scientific Purposes. All experiments were approved by the Norwegian Food Safety Authority.

### Electrode implantation and surgery

Tetrodes were constructed from four twisted 17-μm polyimide-coated platinum-iridium (90%/10%) wires (California Fine Wire). The electrode tips were plated with platinum to reduce electrode impedances to between 120–300 kΩ at 1kHz.

Anesthesia was induced by placing the animal in a closed Plexiglas box filled with 5% isoflurane vapor. Subsequently, the animal received a subcutaneous injection of buprenorphine (0.03 mg kg^−1^), atropine (0.05 mg kg^−1^) and meloxicam (1.0 mg kg^−1^) and was mounted on a stereotactic frame. The animal’s body rested on a heat blanket to maintain its core body temperature during the surgical procedure. Anesthesia was maintained with isoflurane, with air flow at 1.0 liters/min and isoflurane concentration 0.75–3%, as determined according to breathing patterns and reflex responses.

The scalp midline was subcutaneously injected with the local anesthetic lidocaine (0.5%) prior to incision. After removal of the periost, holes were drilled vertically in the skull, into which screws (M1.4) were inserted. Two screws positioned over the cerebellum were used as the electrical ground. Craniotomies were drilled anterior to the transverse sinus. Subsequently, the animal was implanted with either a hyperdrive containing 14 independently moveable tetrodes (seven animals), or a microdrive, containing a single bundle of eight tetrodes. Hyperdrive implants were always on the left side. Hyperdrive tetrodes were implanted perpendicular to the long axis of the HPC (at a 45 deg angle) between −5.5 to −5.7 mm ML and 1.2 to 1.8 AP. Each tetrode was immediately advanced by 940 μm. Microdrive tetrodes were inserted in the right HPC at − 6.9 AP (from right sinus) and 2.0 ML. The tetrodes were immediately inserted to a depth of 1.5 mm. Implants were secured with dental cement (Meliodent). 8–12h after the beginning of the surgery, the animal was treated with an additional dose of buprenorphine (0.03mg kg^−1^).

### Recording procedures

Over the course of 1–3 weeks, tetrodes were lowered in steps of 320μm or less, until high-amplitude theta-modulated activity appeared in the local field potential at a depth of approximately 2.0mm. In hyperdrive experiments, at least one of the tetrodes was used to record a reference signal from white matter areas. The drive was connected to a multichannel, impedance matching, unity gain headstage. The output of the headstage was conducted via a lightweight multiwire tether cable and through a slip-ring commutator to a Neuralynx data acquisition system (Neuralynx, Tucson, AZ; Neuralynx Digital Lynx SX, for all hyperdrive-implanted animals) or via a counterbalanced lightweight multiwire cable to an Axona acquisition system (Axona Ltd., Herts, U.K., for the one microdrive-implanted animal). Both cables allowed the animal to move freely within the available space. Unit activity was amplified by a factor of 3,000–5,000 and bandpass filtered 600–6,000Hz (Neuralynx) or 800–6,700Hz (Axona). Spike waveforms above a threshold set by the experimenter (50–80 μV) were timestamped and digitized at 32kHz (Neuralynx) or 48kHz (Axona) for 1ms. In some Neuralynx recordings, the raw signals were also recorded (32kHz). Local field potential (LFP) signals were recorded from one per tetrode for the hyperdrives and one in total per Axona microdrive. LFP signals were amplified by a factor of 250–1,000, low-pass filtered at 300–475Hz and sampled at 1,800– 2,500Hz. The LFP channels were recorded referenced to the ground screw positioned above the animal’s cerebellum (Neuralynx and Neuropixels) or against an electrode from one microdrive tetrode (Axona). For Neuralynx and Axona recordings, LEDs on the headstage were used to track the animal’s movements at a sampling rate of 25Hz (Neuralynx) or 50Hz (Axona).

### Behavioral procedures

The rats were food restricted, maintaining their weight at a minimum of 90% of their free-feeding body weight, and were food deprived 12–16h before each training or recording session. During the 3–6 weeks prior to surgery and testing, the animals were trained to find the wells filled with chocolate oat milk and alternate between targeted search and direct run towards the home well (ST). When animals achieved a success rate higher than 90% in the ST they were also introduced to random foraging in the open field (OF) (approximately 1 to 1.5 weeks prior to implantation). The arena was a 150×150cm square box with a black floor mat during the OF and a rubber-spray covered plastic plate with 37 small hemisphere incisions of 1 cm diameter arranged in a regular lattice. A cue card (a white A4 paper) on one of the walls indicated orientation. The two environments were located in the exact same place with all distal cues constant and surrounded by the same 50-cm-high black walls. As a reward, vanilla or chocolate biscuit crumbs were randomly scattered in the OF condition and in the ST, chocolate oat milk was provided via the wells in the floor. Curtains were not used, and abundant visual cues were available to the foraging rat. Between sessions in the OF and ST, the rat was placed next to the arena on an elevated flowerpot lined with towels.

Each recording day consisted of two OF and two ST sessions of each roughly 30 min. The ST was run such that all 36 random wells had to be visited at least once, resulting in 36 to 38 home run trials. Care was given that all the arena floors and walls were clean prior to beginning each recording. Data from multiple sessions of the same type was concatenated for analysis purposes. Recordings were generally performed during the dark phase of the 12h/12h light cycle.

### Histology and reconstruction of tetrode placement

Rats where anesthetized with isoflurane (5%) and then received an overdose of sodium pentobarbital. They were subsequently perfused intracardially with saline followed by 4% formaldehyde. The brains were extracted, stored in 4% formaldehyde to be later frozen and cut in coronal sections (animals 20360, 22295, 23783, 24101 and 24116) and para-coronal sections (animal 20382, 21012 and 22098; sections were 45 degrees offset of coronal and sagittal sections in order to align with the angle of drive implantation). Sections of 30μm were subsequently stained with cresyl violet (Nissl) and the relevant parts of CA1 and SUB where collected for analysis. For hyperdrive implants, all tetrodes from the 14-tetrode bundle were individually identified from digital photomicrographs by comparing tetrode traces from successive sections.

The depth of the recording sites was determined post hoc by comparing the deepest visible electrode trace in the tissue with the distance the electrodes were moved between the recording day and the day the animal was perfused. E.g. in Figure 1B tetrode 5 is visible in two images: the leftmost and the middle image in the second row. The track in the middle image, indicated with an empty red triangle (second triangle from the left), is the lowest point in the brain sections where the electrode track could be observed. Between the day the animal was perfused and the last day a neuron had been recorded on tetrode no. 5 the electrode had been moved 270 μm. We therefore assume that the recording sites for this electrode had been approximately 270 μm above this point. This is located in the proximal SUB which at this proximo-distal level extends above the CA1 cell layer. For tetrode no. 12 in animal 20382 the electrode was 100 μm above the recording end point and in animal 20360 the last recordings of tetrodes 8, 9 and 11 have been 100, 210 and 300 μm above the location of the electrode-tip on the day of perfusion. For four-tetrode microdrives, the tetrode bundle as a whole was localized with a similar method.

In order to approximate the anatomical distribution of the recorded cells along the proximo-distal axis, the respective regions were subdivided into proximal, middle and distal CA1 and proximal or distal SUB, respectively. The boundary between the respective subregions was approximated by dividing the region in every section into three or two (for CA1 or SUB, respectively) equally large subregions (see black dashed lines in sections of supplementary Figure 1).

### Data analysis and statistics

Data analyses were performed with custom-written scripts in MATLAB (MathWorks). All statistical tests were two tailed. Each test used was judged to be the most suitable for the data in question. Where possible, nonparametric methods were used in preference to parametric methods, to avoid making unwarranted assumptions about the underlying data distributions. The study did not involve any experimental subject groups; therefore, random allocation and experimenter blinding did not apply and were not performed.

### Spike sorting and single-unit selection

Spike sorting was performed offline using manual cluster cutting methods with MClust (A. D. Redish, http://redishlab.neuroscience.umn.edu/MClust/MClust.html). Spike rate autocorrelation and cross-correlation were used as additional tools for separating or merging spike clusters. Single units were discarded if more than 0.5% of their inter-spike interval distribution was comprised of intervals less than 2ms or if they had a mean spike rate less than 0.5Hz. Additionally, interneurons were separated and excluded from the dataset by removing clusters with narrow waveforms and high firing rates during manual spike sorting.

### Behavioral parameters extraction

Animal position was estimated by tracking the LEDs or reflective markers mounted on the implant. Only time epochs in which the animal was moving at a speed above 2cms^−1^ were used for spatial analyses.

To generate 2D rate maps for the open-field arena, position estimates were binned into a 5×5cm square grid. The spike rate in each position bin was calculated as the number of spikes recorded in the bin, divided by the time the animal spent in the bin. The resultant 2D rate map was smoothed with a Gaussian kernel with σ=1.5 bins.

The animal’s head direction was determined from the relative positions of LEDs or reflective markers on the implant. Head direction tuning curves were calculated by binning the head direction estimates into 6° bins. The spike rate in each angular bin was calculated as the number of spikes recorded in the bin divided by the time the animal spent in the bin. The resultant tuning curve was smoothed with a Gaussian kernel with σ=1 bin, with the ends of the tuning curve wrapped together.

### Spatial correlation across session type

For the spatial correlations across session types (i.e. spatial memory task (ST) vs open field (OF) the data from open field and foster maze were concatenated, respectively. Half of the session was chosen by taking 5 minutes intervals of the data which then were shifted forward n times with 30 second intervals. For each shifted sub-portion of the data, the rate map was calculated and correlated with the other half of the data. The number of shifts needed to get a stable value was determined in a saturation process where for every cell increasing number of shifts were used until three consecutive values of the correlation were below 0.5 standard deviations.

### Calculating information rate and information content

Information rate (***I***_***rate***_) and information content (***I***_***content***_) were calculated as described in Skaggs et al. (1993). Briefly, every covariate space was binned (900 bins of 5 × 5 cm for position, 10 bins for head direction and 10 bins for speed) and the neurons spikes were allocated to the respective bin the animal was occupying while they were emitted. The total number of spikes in every bin was then divided by the total session time to receive ***λ(x)***, the average firing rate in bin ***x***. Using ***λ***, the mean firing rate of the neuron across the entire session and ***P(x)***, the probability of the animal to occupy bin ***x***, information rate for each neuron across the entire session was calculated by:

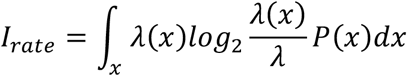

To calculate information content per spike, ***I***_***rate***_ was divided by the average firing rate of the neuron across the entire session ***λ***.

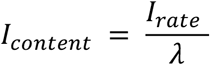

In order to ensure that the information values are minimally affected by any biases, we subtracted from all information measures the mean of the value obtained for the same measure for 100 shuffled spike distributions. The values reported in the text are with these biases removed, although our results do not qualitatively change if we do not remove the bias.

### Spatial fields extraction (for border score and boundary vector score calculations)

For extracting spatial fields from the rate map of the neurons we used two thresholds: one for detecting and defining the borders of the field and a second one for including the field. First all the pixels inside a field have to be higher than the median plus one standard deviation of the map (**field detection threshold**). Then any valid field had to have a peak firing rate equal or higher than the median plus two standard deviations of the map (**field inclusion threshold**). The fields were extracted with the Matlab ‘contour’-function and the field detection threshold was applied with ‘inpolygon’. All the fields with a peak firing rate higher than the field inclusion threshold were concatenated to a single so-called fields-map.

### Preparation of the data for the Poisson GLM

Spike time stamps obtained from the spike sorting procedure were binned using bins of length ***dt*** = 20 ***ms*** to get a spike count train vector 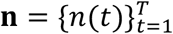 per each cell, where ***n(t)*** is the spike count in time bin ***t***, while ***T*** is the total number of time bins. The behavioral covariates taken into account are:

P: xy-position of the rat on the horizontal plane;
H: head direction of the rat on the horizontal plane;
S: running speed of the rat;

while the internal covariates consist of:

T: theta phase, i.e. the phase of the band-pass filtered (5 to 12 Hz) and Hilbert-transformed local field potential.
E: ensemble spike signal, defined as the z-scored sum of the spike counts of all neurons simultaneously recorded at the same tetrodes.

All covariates were interpolated at the centers of the time bins to achieve the same temporal resolution of the spike counts. By means of an optimization procedure over the entire population of neurons (see section Hyperparameters optimization) we chose to bin each covariate ***C*** in to ***N_C_*** bins. Finally for each covariate ***C*** in each time bin ***t***, we built a binary state vector ***X^C^(t)*** of length ***N_C_***, with entries 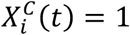 if the animal behavioral state at time ***t*** fell in the ***i***-th bin for the covariate ***C***, while 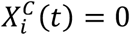 otherwise. Unless explicitly stated the OF and ST sessions were concatenated.

### Poisson GLM

We adopted a Poisson GLM to explicitly model the stochastic response of each neuron to the covariates ***C***. The choice of a Poisson random component as used in Hardcastle et al. (2017) is further motivated in subsection ‘*Poissonian spiking*’. The models 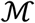 we consider, are combinations of behavioral and internal covariates ***C* = *P, H, S, T, E***. For a given model 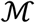 the probability of recording ***k*** spikes in time bin ***t*** of length ***dt*** is Poissonian:

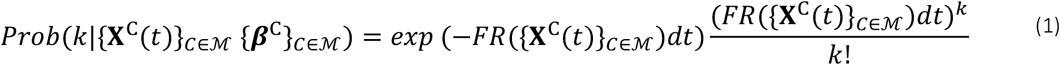

where 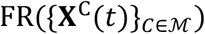 is the expected firing rate in time bin ***t***, and

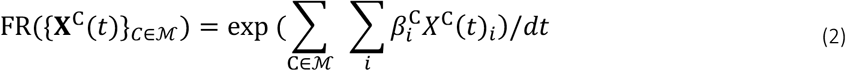

depends on the behavioral/internal state vectors ***X***^***C***^ in time bin ***t*** and the vector of predictors ***β***^***C***^ for all covariates 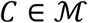. The predictors are estimated by the learning procedure explained in section ‘*Learning*’.

#### Poisson spiking

We selected the Poisson distribution to fit the stochasticity of the firing process after explorative analysis of our data and rigorous testing. First, we observed that the spiking data are not binary for time bin lengths ***dt ≥ 1ms***. This indicated that the Bernoulli distribution may not be a suitable choice. Secondly, we verified that the inter-spike-interval (ISI) distribution is well fitted by an exponential up to 60 ms (the deviations at 1-2 ms can be attributed to the refractory period). The ISI distribution expected for Poisson firing is indeed exponential.

In addition, we looked at the Fano Factors of the spike count ***N*** for each cell, defined as:

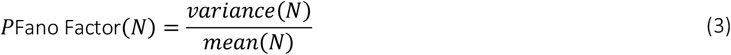

This analysis indicated that the ***variance(N)*** versus ***mean(N)*** plots are well fitted by the line that bisects the first quadrant of the coordinate system for time bin lengths ***dt ≤ 40ms***. The coefficient of determination of the fit ***R*** – ***square*** is an indication of how much the spiking process deviates from a homogeneous Poisson process, for which a Fano Factor of 1 is expected in the long run.

Notice that the Poisson GLM in Equation (1) is not a homogeneous Poisson process (the firing rate of Equation (1) depends on the covariates which vary in time), it becomes homogenous Poisson once the covariates in Equation (1) are fixed. We used the asymptotic distribution of the Fano Factor for the homogeneous Poisson (Eden and Kramer, 2010) to test how poissonian the spike counts are at fixed covariates. We run the test after conditioning the spike counts on the covariate position in a 30 times 30 grid superimposed to the recording box. The number of position bins was optimized for this test and bins with occupancy shorter than 250 ms were excluded by this analysis. The test with 5% significance level resulted in an average fraction of position bins rejected by cell of 0.0589 ±0.0009. Finally, instead of conditioning on position, we conditioned on time, by testing the Fano Factor of the spike count in 500 ms time windows. The conditioning in time is motivated by slow varying covariates which enforce to the firing rate a different time scale from the spiking time scale. This approach resulted into an even smaller average rejection rate, suggesting that position may not be the only covariate modulating the activity of this cells. The average fraction of 500 ms time windows in which the Poisson hypothesis has to be rejected with a 5% confidence is 0.0152 ±0.0002.

### Encoding

#### Learning

In order to determine the selectivity of a neuron to the covariates ***C = P, H, S, T, E***, given the recorded spike count train vector **k** and the vectors of covariates ***X^C^(t)*** in each time bin ***t***, we optimized the predictors 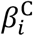 of the Poisson GLM in Equation (1) for each model 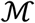 to minimize the cost function:

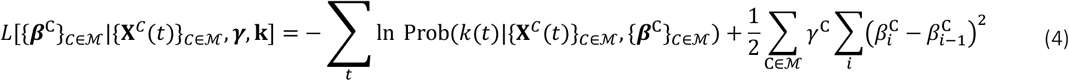

such that the learned parameters are 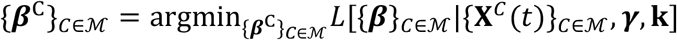. The first term in the loss function of Equation (4) is the negative log-likelihood of the spike count train vector **k**, while the second term is a penalty on large differences in parameters between nearest neighboring covariate bins and therefore enforces smoothness in the model predicted tuning curves. The smoothness hyperparameter ***γ***^***C***^ controls the strength of the smoothness penalty for the covariate **C** and was optimized a priori on the entire population of neurons, as explained in section Hyperparameters optimization. The minimization of the function 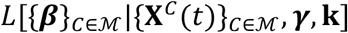 in the parameter space was performed using the Matlab *minunc* function. The learned parameters ***β*** were used for estimating model performance, constructing model predicted tuning curves (see section Tuning curves predicted by the model) and decoding.

In addition to the models containing the behavioral/internal covariates we also fitted a model with only the bias term **(FR(*β*_0_) = ex p(*β*_0_) /*dt*)**, which we term the average firing rate model. Model performance for each cell was assessed in a 10-fold cross-validation setup: of the whole spike count train one tenth was held out as test set, while the rest constituted the training dataset, on which parameters were learned. The difference in log-likelihood of the test spike count train between the model 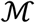 and the average firing rate Poisson model (also learned on the training data) was taken and then divided by the number of time bins in the test set. The cross-validation procedure was repeated for 10 non-overlapping test sets, resulting in 10 values of the log-likelihood increase over the average firing rate model per time bin 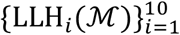. Their average across cross-validation folds 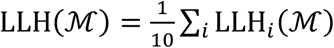 was regarded as the main indicator of model 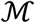 performance.

#### Hyperparameters optimization

For optimizing the hyperparameters, number of bins ***N_C_*** and smoothness hyperparameter ***γ***^***C***^ for each covariate ***C = P, H, S, T, E***, we computed the model performance **LLH(C)** on a grid in the hyperparameters space (***N_C_* = {2,5,10,20,30,40,50,60,70}, *γ^C^* = {0.08, 0.8,0,80,800})**, for each of the single covariate model 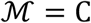 separately and for each cell independently. The values of **LLH(C)** at fixed hyperparameters were then averaged over all cells in the population, and the values of the hyperparameters at its maximum on the grid were taken as candidate optimal hyperparameters. To compare model performance at the maximum to its neighboring vertices on the grid in the hyperparameters space, we employed the one tailed Wilcoxon signed-rank test between the population vector of **LLH(C)** at the maximum and the same vector in its nearest neighbors on the grid. The values of the hyperparameters including the smallest number of covariate bins ***N_C_***, whose corresponding vector of average normalized log-likelihoods was not significantly different from the maximum (p > 0.05 in the Wilcoxon signed-rank test), were chosen as new candidate optimal hyperparameters. This procedure was then iterated until all neighboring models with smaller numbers of covariate bins were significantly different in model performance according to our test. The hyperparameters were then held fixed during learning on the entire population of neurons and for all models.

The optimal number of bins and smoothness hyperparameters are respectively ***N_P_* = 30** (along each edge of the squared enclosure, 900 in total) and ***γ^P^* = 8** for position, ***N_H_* = 10** and ***γ^H^* = 800** for head direction, ***N_P_* = 10** and ***γ^S^* = 800** for speed, ***N_T_* = 10** and ***γ^T^* = 800** for theta, ***N_E_* = 20** and ***γ^E^* = 80** for ensemble activity.

#### Model selection

Learning was performed for all 31 models 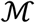, i.e. the single covariate models ***P, H, S, E, T***, the two covariate models ***PH, PS, HS, PE, PT***, …, all the three, the four covariate models up to the five covariates model ***PHSET***. The learning method is described in section Learning. We adopted a forward model selection procedure (Hardcastle et al., 2017) aiming at selecting the model with the highest performance and the smallest number of covariates. As a starting point we used the average firing rate model (only bias). So, we considered the model with the largest performance 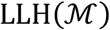 among the single covariate models, e.g. 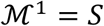. The vector of log-likelihood increases over the average firing rate model 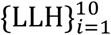 across cross-validation folds for the selected single covariate model 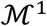 was compared to the same vector for the average firing rate model (a vector of zeros), by means of the one-tailed Wilcoxon signed rank test. If significantly different (***p < α*** in the one tailed Wilcoxon signed rank test), then the single covariate model became the new candidate selected model, otherwise the average firing rate model was selected and the search in the model space interrupted. In the former case, the model with the largest 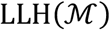 among the two covariate models, including the covariate selected in the former step, e.g. 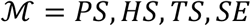, was considered, e.g. 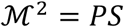. In full analogy with the previous selection step, the vector of log-likelihood increases over the average firing rate model 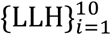 across cross-validation folds for the single covariate model 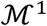 was compared to the same vector for the two covariate model 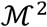, by means of the Wilcoxon signed-rank test. If the mean of the vector for 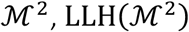, was significantly larger than the mean of the vector for 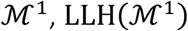,(***p < α*** in the one tailed Wilcoxon signed-rank test) the model 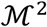 was identified as candidate model. If not significant the search in the model space was interrupted and the single covariate model 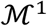 became the selected model. The forward search was carried on in this fashion by including one covariate at each step to the candidate selected model, up to the selected model resulting in a not significantly different outcome in the Wilcoxon signed-rank test. We checked the sensitivity of the model selection procedure to the significance level ***α***.

#### Splitting between external and brain internal variables

In the current study we were interested in how CA1 and SUB neurons compare in their mapping of navigational covariates P, H and S. The covariates E and T (ensemble firing and theta) were treated as brain internal states which helped fitting the model but did not directly explain the behavior of the animal. We therefore used them as auxiliary variables which helped to get the best possible fit, by on the one hand 'explaining away' spikes that otherwise would have been attributed to position, head direction or speed and on the other reducing the level of noise in the spike trains such that other spikes could be better attributed to the correct navigational covariate. For example, it might be conceivable that a neuron has a higher firing rate always when the the other neurons at the tetrodes (E) have a high firing rate because the neuron is not well enough isolated from the ensemble. In a scenario in which the ensemble responds to speed, including E in our model will attribute some spikes of that neuron to ensemble firing and less to the speed response. For an extended graph with all models displayed separately, see Figure S4H.

#### Relative single covariate contributions (rSCC)

We measured the contribution of a single covariate **C** included in the selected model 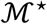 in explaining the firing rate of a neuron in terms of the difference in model performance between the selected model and the model resulting from removing the covariate **C** from the selected model, 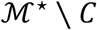. We therefore defined the rSCC (**C**) for covariate 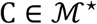 as 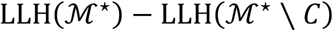 normalized by

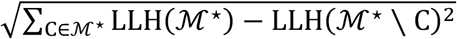

where 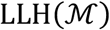 is the normalized log-likelihood of the model 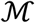 on the held-out data averaged across cross-validation folds of ***PHS, C*** refers to the two-covariate model that does not include the covariate ***C***. Single covariate contributions of covariates not included in the selected model were set to zero.

#### Mixed selectivity score (MS-score)

The MS-score was defined as the product of rSCC of position, head direction and speed:

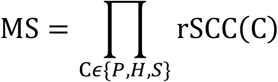

#### Tuning curves predicted by the model

Model predicted tuning curves were constructed on the basis of the model including all covariates, 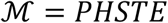, whose parameters ***β*** were learned as explained in section ‘Learning’. A tuning curve is the average firing rate of a neuron as a function of relevant stimulus parameters, in our case covariate bins; for the model predicted tuning curves the average is taken over the distribution of the other covariates. In case of uniform sampling and assuming independence between covariates, the expected value of the firing rate in the ***i***-th bin of covariate **C**^∗^ for Poisson neurons as defined in Equation (4) is:

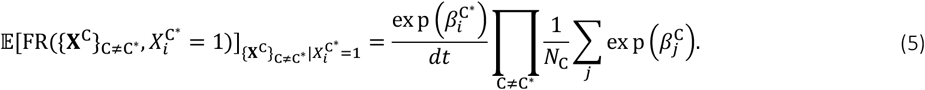

#### Goodness of fit

In order to assess the goodness of fit of the GLM, in addition to 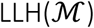, the average of the explained deviance across cross validation folds 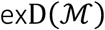 was estimated (Brown et al., 2002; Guisan and Zimmermann, 2000; Kraus et al., 2015; Pillow, 2009). The explained deviance of model 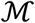 in cross-validation fold i is defined as 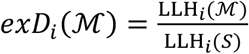 where S is the so-called saturated model, a maximum-likelihood Poisson GLM optimized on the test data (in practice the firing rate FR in Eq. (2) is set equal to the spike count k(t) in each time bin t). Then 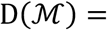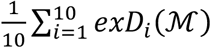 quantifies the fraction ascribable to model 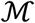 of log-likelihood increase with respect to the average firing rate model of the best Poisson model. For Figure S11 the goodness of fit of the GLM as a model was evaluated for different region and tasks, therefore per each cell only the best fitting model 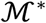 was been considered.

#### Testing for significance in model selection between CA1 or SUB and between OF or ST

For Figures 4E and 5B we used a shuffling procedure to test whether the type of model selected for a proportion of neurons was significantly affected by the anatomical location of the neurons or the task in which the neurons were recorded. For the pair of conditions being compared (either CA1 vs. SUB, or OF vs. ST), each neuron was randomly assigned to one of the two conditions with equal probability, and for each model (e.g. PH or PHS) the absolute difference was calculated between the proportion of neurons selecting it in the two conditions. This process was repeated 10000 times, yielding a distribution of shuffled differences for each model. If the actual difference between the two groups was outside the 99.9^th^ percentile of 10000 such shuffled differences, we considered the difference for a given model (e.g. PH or PHS) to be significant.

### Decoding

We exploited the Poisson GLM framework defined in section Poisson GLM for decoding simultaneously position P, head direction H and speed S. We employed the model including all behavioral covariates ***P, H, S*** and the internal covariate theta T. Given the learned parameters ***β***^***c***^ with ***C = P, H, S ∪ I***, the vectors of spike counts **k_n_**, for cells ***n* = 1, …, *N*** and the internal observed covariates ***X^I^*(*t*)**, the decoded covariates at time ***t***, {***X*^C^(*t*)}_*C∈PHS*_**, were chosen as those maximizing the objective function:

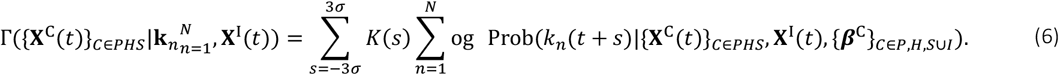

In Equation (6) *K*(*S*)is a gaussian kernel of zero mean and standard deviation *σ*, while **Prob(*k*_n_(*t* + *S*)|{*X*^C^(*t*)}_*C∈PHS*_, *X*^I^(*t*), {*β*^C^}_*C∈P,H,S*∪*I*_)** is the probability of observing the spike count ***k_n_*** at time *t* + *S*, given the covariates{***X^C^*(*t*)}_*C∈PHS*_, *X^I^*(*t*)** according to the Poisson model in Equation (2) with parameters. **{*β^C^*}_*C∈P,H,S*∪*I*_**

Decoding is then maximizing the weighted log-likelihood of the spike counts of all cells in a time window of size 6*σ* + 1 centered at the time point ***t***. The parameter *σ* has been optimized for the entire dataset (both regions and both tasks).

Maximization of the objective function Γ was performed by grid search on the covariates lattice defined by the binning adopted for learning, where the number of bins has been optimized as explained in Section ‘Hyperparameters optimization’.

### Resampling of neurons

For decoding we sampled different sized subpopulations (from 5 to 150 neurons) and maximized the likelihood of the spike counts in the space of all three behavioral covariates. This allowed us to decode the tree covariates P, H and S simultaneously at every time bin t by optimizing the likelihood of the weighted spike counts from Gaussian filtered time window *t* − Δ*t* to *t* + Δ*t* (where 2Δ*t* is the size of the decoding window: 200 ms) assuming that the covariate did not change in this specific time window. We, furthermore, resampled neurons recorded in the OF and in the ST separately, in order to asses if the neural response during attentive behaviors carried a better decodable signal than during random foraging.

#### Randomization of decoding

The randomized decoding distributions for Figure S6, are built by taking the difference between two means from decoding 20 times from a randomly selected subpopulation of neurons, while ensuring equal probability to select CA1 or SUB neurons. The randomization runs over 1000 iterations and it is assessed whether the difference between decoding error of CA1 and SUB population is larger than the 99^th^ percentile of the randomized differences.

### Code availability

Code for reproducing the analyses in this article is available from the corresponding authors upon reasonable request.

## Acknowledgments

We thank A.M.Amundsgård, K.Haugen, E.Kråkvik, H.Waade, and V.Frolov for technical assistance and Hiroshi T. Ito for advice and discussion. The work was supported by the European Commission’s FP7 FET Proactive Programme on Neuro-Bio-Inspired Systems (GRIDMAP, Grant Agreement 600725), two Advanced Investigator Grants from the European Research Council (GRIDCODE, grant no. 338865; ENSEMBLE, grant no. 268598), the Centre of Excellence scheme and the National Infrastructure scheme of the Research Council of Norway (Centre for Neural Computation, grant number 223262; NORBRAIN1, grant number 197467), and the Kavli Foundation.

## Figure Legends: Supplementary Figures and Tables

**Figure S1.**
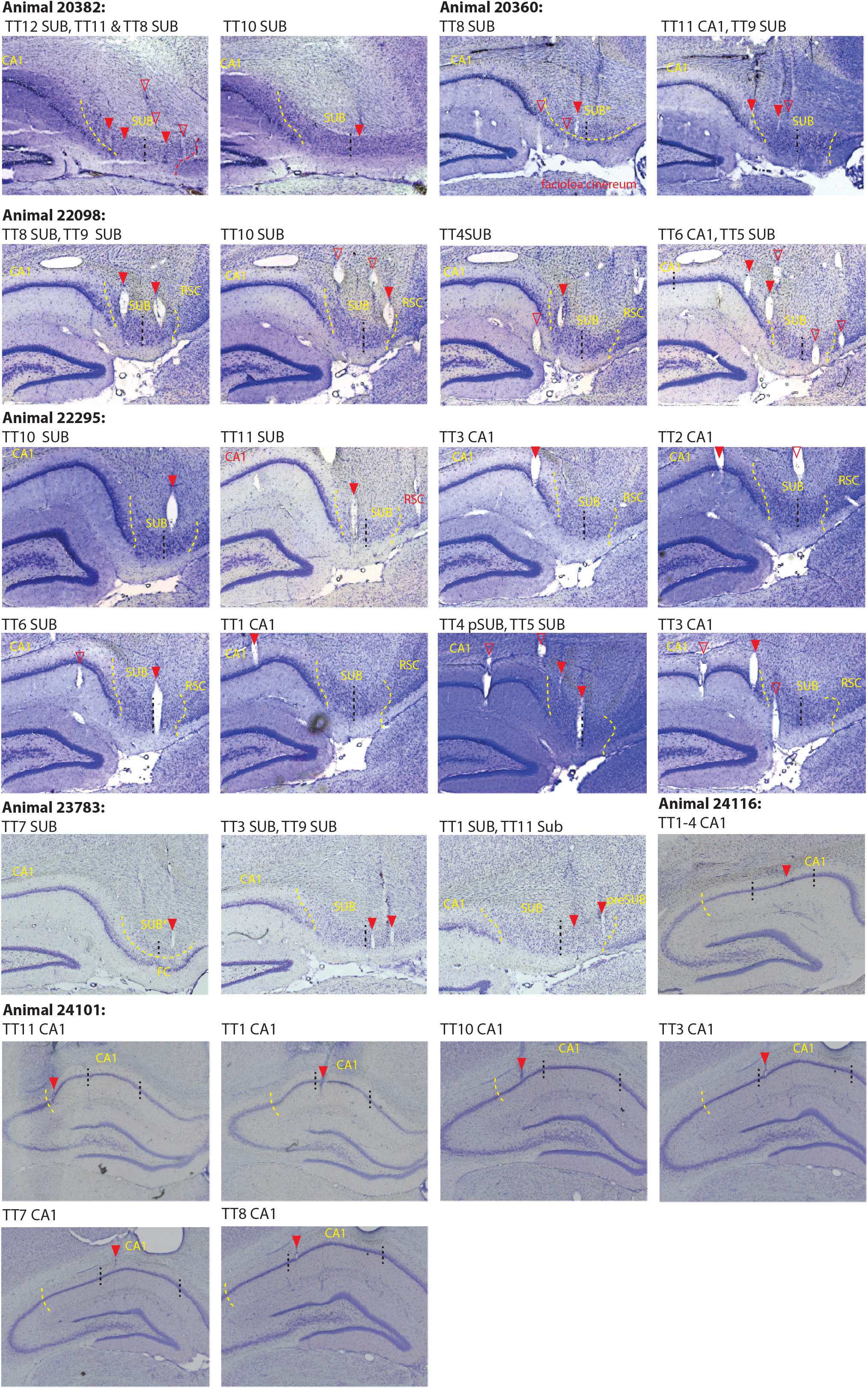
Anatomical locations of recording electrodes. Nissl-stained sections for every animal recorded in the study. Animal ID number listed on top of the Nissl section marks the beginning of sections from a new animal and so applies to all following sections. Filled red arrows indicate tetrode (TT) traces at the estimated recoding location (usually at the end of the tetrode track) while empty red arrowheads indicate traces of tetrodes that are visible in this section but are above (or below) the locations on the track where the recordings were conducted. The TT number above each Nissl section indicates the identity of tetrodes with filled arrowheads, arranged from left to right on the section. Dashed yellow lines indicate borders between SUB and neighboring regions like retrosplenial cortex (RSC), presubiculum (preSUB) and faciola cinerata (FC). For recording locations in SUB that are dorsal to the FC (labeled with SUB*) or the border between CA1 and SUB, special care was given to include data only from neurons that could confidently be assigned to be 50 μm or more away from the border.

**Figure S2.**
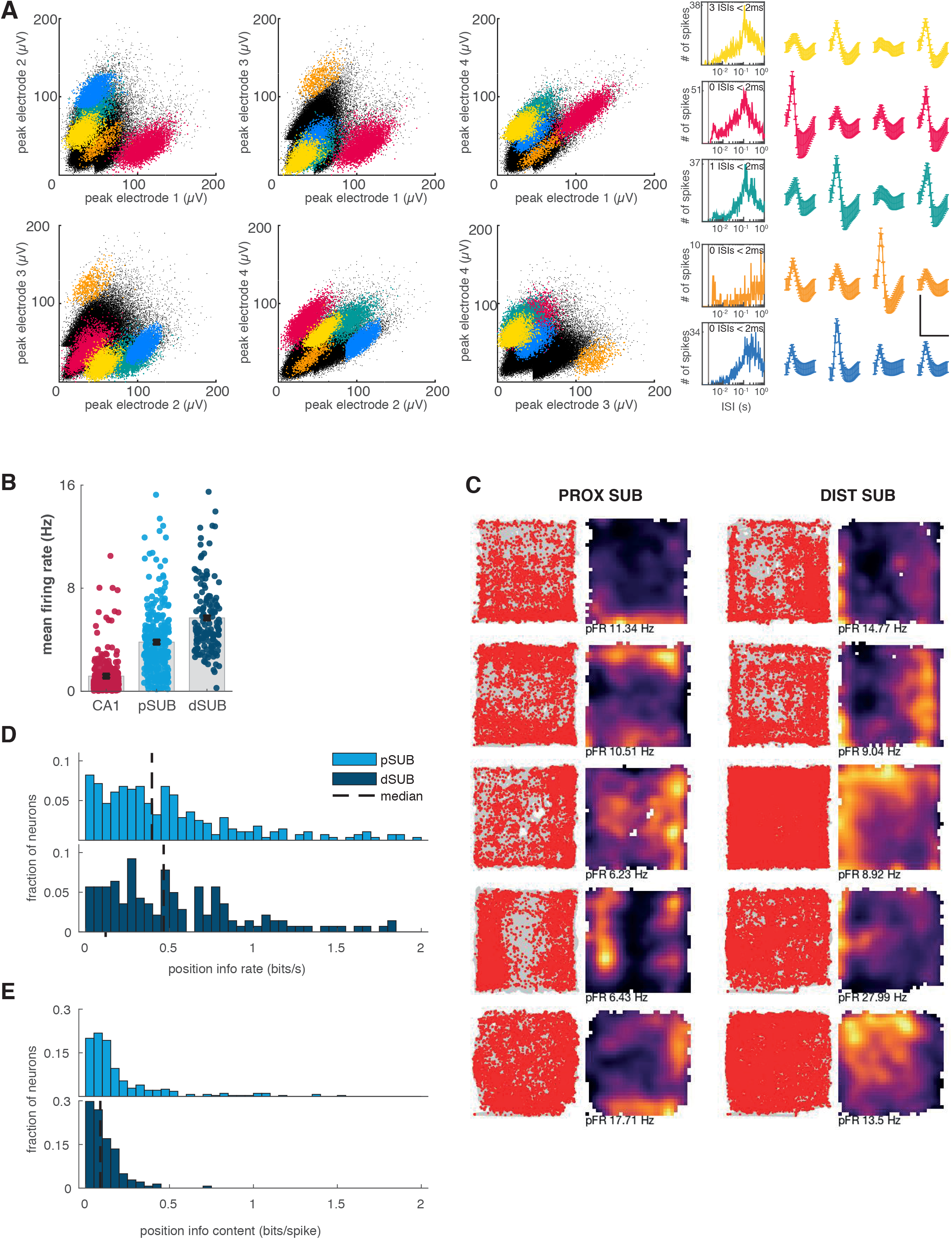
Spatial firing fields of principal neurons along the proximo-distal axis of the SUB. (A) Multidimensional cluster diagrams illustrating isolation of spikes from simultaneously recorded cells in SUB (the yellow cluster is the same cell as shown in the first, left panel of Figure S2C). Left: Scatter plots showing relationship between peak-to-peak amplitudes of spikes recorded from electrodes of a tetrode. Each point represents one sampled signal. Clusters are likely to correspond to spikes originating from the same cell. Distinct clusters are assigned unique colors. Middle: spike autocorrelograms showing spike frequency as a function of interspike interval (ISI) for each cluster isolated in the scatterplot. Note absence of spikes at intervals less than 2 milliseconds (gray vertical line), as expected due to refractory period for spikes originating from the same cell (number of spikes with ISI < 2 ms is indicated for each unit). Interval scale is logarithmic. Right: Waveforms on four channels for the cells isolated in the scatterplot (means ± standard deviations; scale bars: 100 μV, 1 ms; colors as in the scatter plot). (B) Mean firing rates of all principal neurons recorded in CA1 (red), proximal SUB (light blue) and distal SUB (dark blue). Mean values (+-S.E.M.) of all neurons are shown in black and by the edge of the gray bar in the background. The average firing rate increased significantly from CA1 1.2±0.1 to SUB 4.4±0.1 and within SUB from 3.8 ±0.2 Hz in the proximal half of SUB to 5.7 ±0.2 Hz in the distal half (p = 5.1e-11, Welch’s test; Figures S2A left panel and S1B). (C) Representative path plots (left) and rate maps (right) from neurons in the proximal (left column) and the distal SUB. Rate maps show firing rates color-coded from 0 (black) to the neuron’s peak firing rate (pFR – white), indicated below each rate map. Color scale as in Fig. 2A. Path plots show the path of the animal during the entire session in gray and the emitted spikes of the respective neuron overlaid in red. (D-E) Frequency distributions showing position information rare (D) and information content (E) for all neurons in proximal (light blue) and distal SUB (dark blue). The dashed line indicates the 99^th^ percentile of a distribution of shuffled data. The percentage of neurons passing the 99^th^ percentile of the shuffled distribution for position information content in C is 7.9% in pSUB and 0.7 % in dSUB. For position information rate (D) the percentage is 22.1% in pSUB and 29.8% in dSUB.

**Figure S3.**
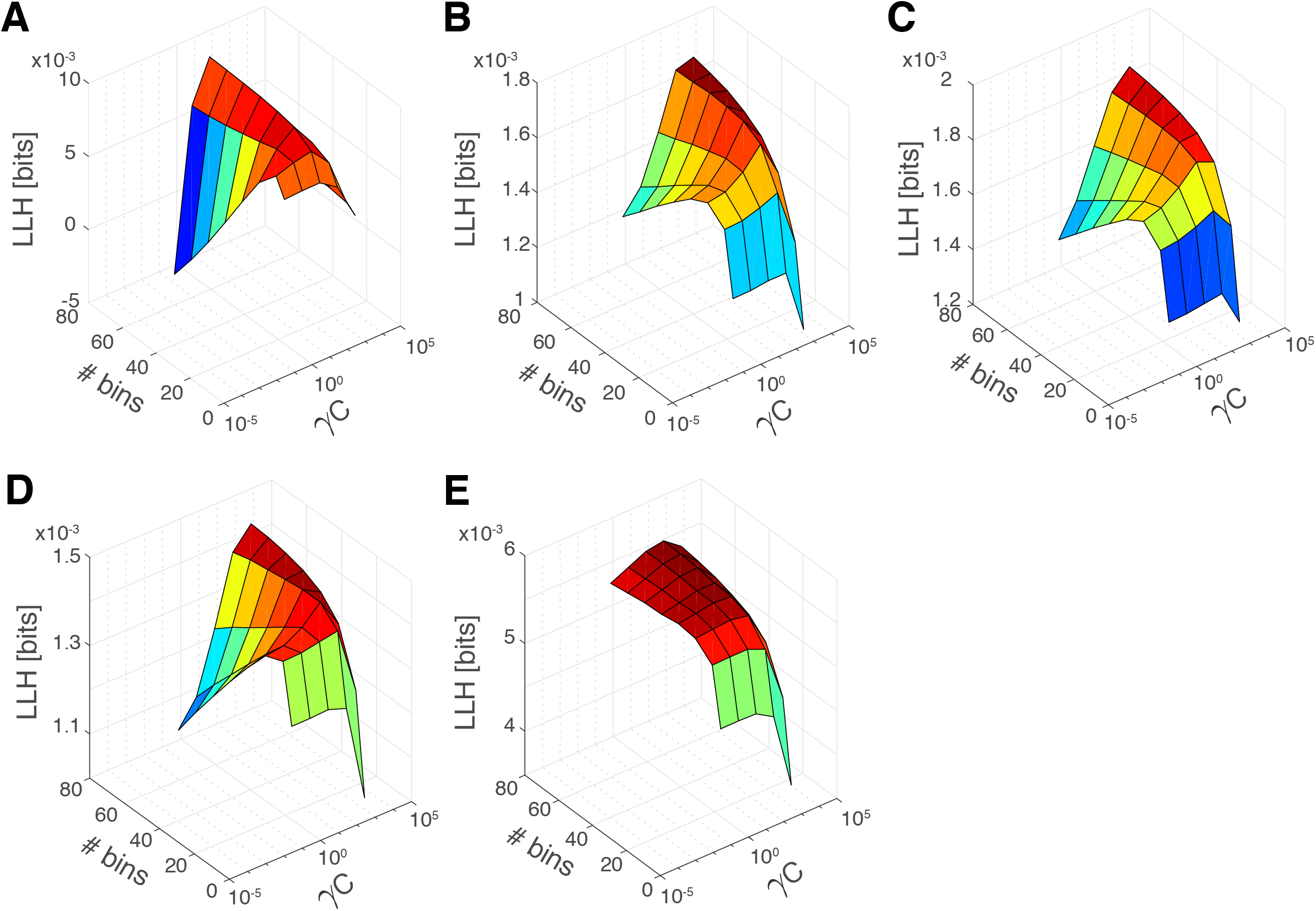
Optimization of hyperparameters. (A-E) Model performance (LLH, see section *‘Learning’* in Methods) averaged across all 746 cells (325 from CA1 and 421 from SUB) for position (A), head direction (B), speed (C), ensemble firing (D) and theta phase (E). Single covariates GLMs are plotted as a function of the GLM hyperparameters: number of bins (# bins) and strength of the regularizer (γ_C_). Given the maximum of the model performance, values for hyperparameters were optimized by minimizing the number of parameters without causing a significant decrease of model performance with respect to the maximum. For details, see section *‘Hyperparameters tuning’* in Methods. Blue colors indicate the lowest value and red colors indicate the largest value of the z-Axis (LLH). The bins of the surfaces are colored according to the lowest point on the z-axis that the bin touches. The range of the color-scale is normalized for every figure separately.

**Figure S4.**
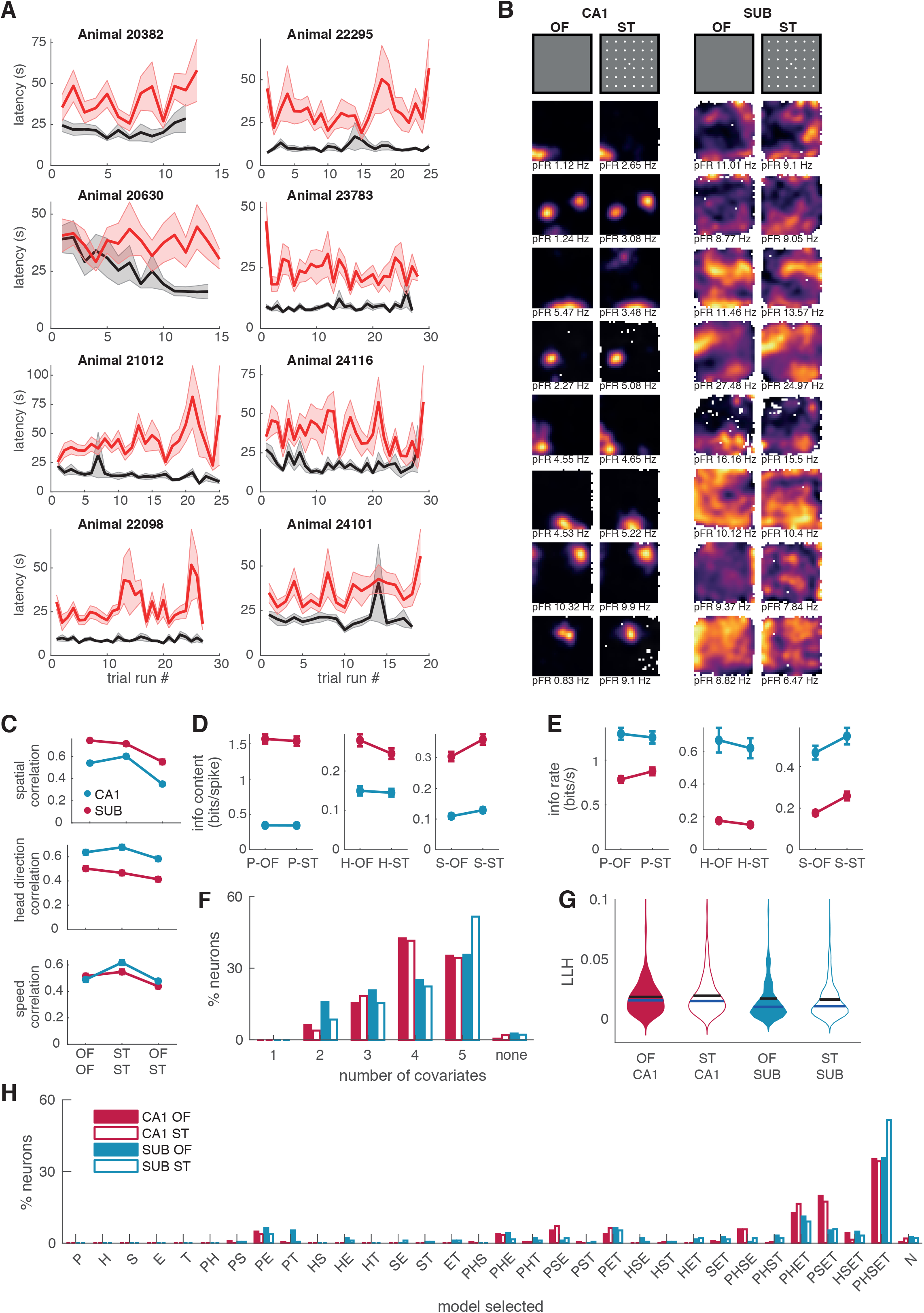
Quantification of neural tuning and behavior in the ST and the OF. (A) Latencies to find the home well (gray) and the random well (red) for eight different rats across all trials of all sessions. Note that for animals 20382, 22295 and 20630 the home well was in a different location in every session while for the other animals it was always in the center of the environment. For these three animals therefore, home run trials have a longer latency at the beginning than at the end of the sessions while for the other animals this latency was low from the start of every session. (B) Rate maps of example cells from the CA1 (left column) and the SUB (right column) in the OF (left) and ST (right). White color indicates peak firing rate, indicated also at the bottom of each map, while black colors indicate zero Hz firing rate (color bar as in Figure S2). (C) Correlations of tuning curves for position, head direction and speed between OF-OF, ST-ST and OF-ST for position (top), head direction (middle) and speed (bottom). Correlations were calculated for every neuron between all pairs of OF and ST sessions the neuron was recorded in. The average correlation of all neurons was then compared within and across session types. Position correlations within and across session types were significantly higher for CA1 than for SUB. For both regions, correlations between same-task sessions were larger than between sessions of different task types (Welch’s test for OF-OF vs. OF-ST comparisons in CA1: p = 1.4e^−16^ and SUB: p = 6.3e^−27^; Welch’s test for ST-ST vs. ST-OF comparisons in CA1: p = 1.02e^− 15^ and SUB: p = 8.71e^−30^). Correlations for head direction tuning curves (middle) were significantly higher in SUB than in CA1 and dropped when same-task-correlations were compared to across-task-correlations (Welch’s test for comparing ST to OFST: p = 3.8e^−8^ and for comparing OF to OFST: p = 1.1e^−3^). For speed tuning curves (bottom), correlations within and across task types were similar in CA1 and SUB except for a small but significant increase in correlation of SUB when correlating ST to ST sessions. The average of ST to ST correlations was significantly higher than OF to OF correlations (p = 1.3e-8) and ST-OF correlations (p = 1.5e-11). This might indicate that the speed tuning in ST is more stable in SUB when compared to other tasks and to CA1. In summary tuning curves in CA1 are more stable in terms of position, while in SUB stability is highest in terms of head direction and speed. (D-E) Information content (D) and information rate (E) in OF and ST for position, head direction and speed. (F) Percentage of neurons selecting a 1, 2, 3, 4 or 5 covariates model in CA1 (red bars) and SUB (blue bars) during the OF (full bars) and the ST (empty bars) condition. When taking into account all the covariates, (P, H, S, T and E) a 5-covariate model was selected for a larger number of neurons in SUB than in CA1. For the majority of CA1 neurons, a 4-covariates model was selected in both the OF and the ST. (G) Distributions of log-likelihoods (LLH) in CA1 (red violin plots) and SUB (blue violin plots) during the OF (full violin plots) and the ST (empty violin plots) condition. The mean and the median of the LLH are comparable between CA1 and SUB neurons and between the neurons recorded in the OF and the ST. The GLM fitting process therefore performs similarly in all conditions. (H) Model selection plot across all 32 possible models without collapsing models containing ensemble firing (E) and theta (T) with the models containing only the external covariates position (P), head direction (H) and speed (S). Separating the models also for E and T shows that the most complex model (PHS in Figure 4) which performed best for a majority of neurons in SUB, most often consists of the 5 covariate model including E and T. The model most often selected for CA1 neurons is the PHET model when the models are separated for E and T. Models missing E and T as covariates (like PHSE or PHST) were only rarely selected. Neurons that are completely independent of ensemble firing or theta were extremely rare (see groups P, H, S, PH, PS, HS and PHS).

**Figure S5.**
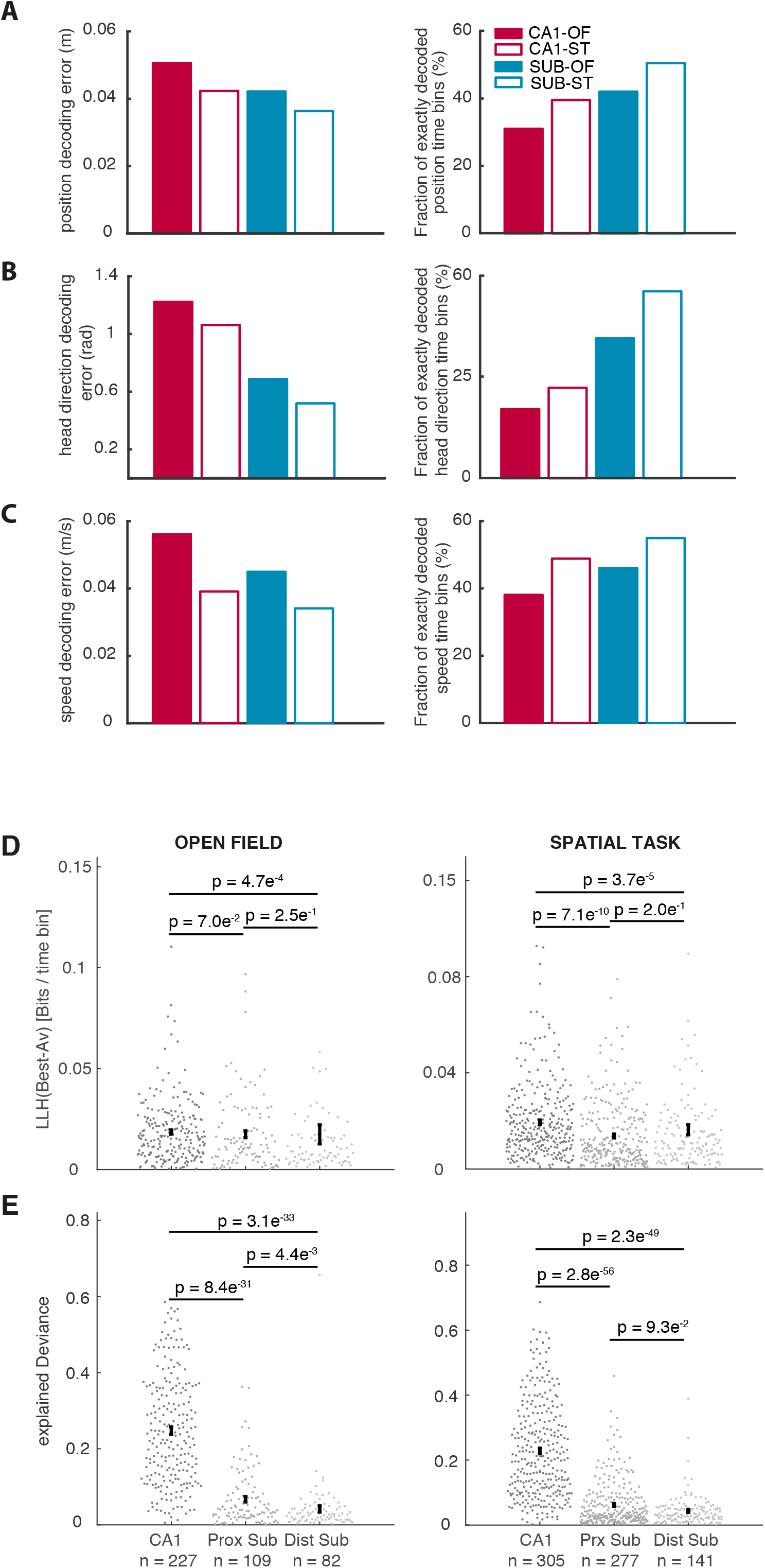
Controls for decoding: Equal sampling from the different bins of speed and goodness-of-fit of the GLM. (A-C) Accurate decoding of position, head direction and speed when the data was equally sampled from the different speed bins. Decoding error (left column) and percentage of exactly decoded bins (right column) at a population size of 150 neurons for position (A), head direction (B) and speed (C). Note that the difference in decoding error for position between OF (full bars) and ST (empty bars) remains after equally sampling from different speed bins between OF and ST. (D-E) Average across all the cross-validation folds of the difference in the log-likelihood per time bin of the best model M* and the average firing rate model – LLH(M*) defined in section Learning of the Methods (D) and cross-validated explained Deviance per time bin – exD(M*) defined in section Learning of the Methods (E) for the same population of neurons recorded in the open field (left column) and the spatial task (right column). Spike trains from neurons in CA1 and SUB are equally well modelled by the GLM by goodness-of-fit measures of explained deviance (D) or difference in likelihood between the best performing and the average firing rate model (E). The superiority of decoding in SUB is therefore unlikely to result from the GLM as a modeling framework. Black dots and error-bars indicate the sample mean and s.e.m., respectively. P-values were computed using the Mann–Whitney U test.

**Figure S6.**
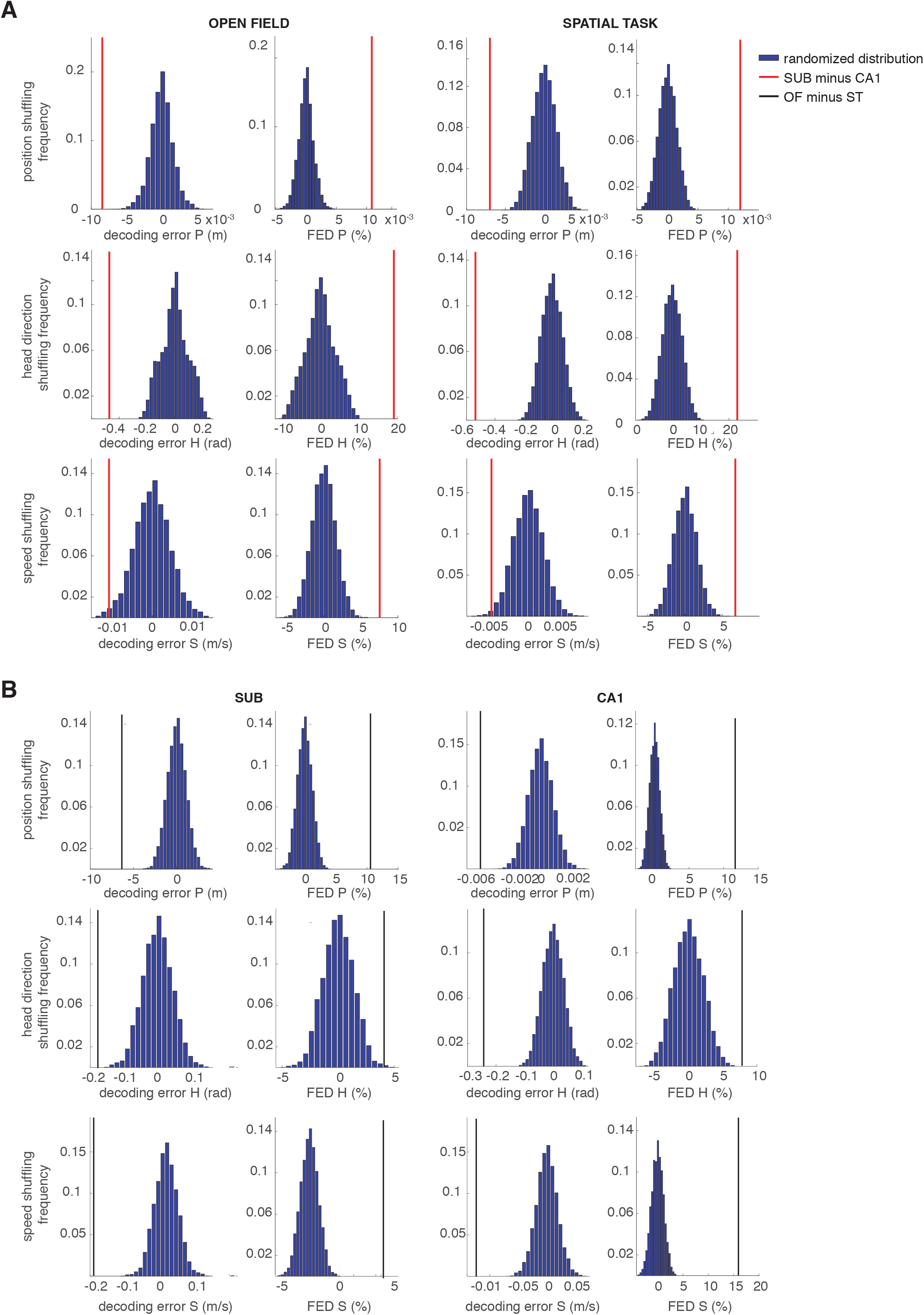
Significant difference in decoding error between SUB and CA1 and between OF and ST when compared to the differences in a shuffled population. (A) The blue histograms show the frequency of occurrence for differences in decoding error (left) and frequency of exactly decoded bins (FED) when decoding from populations of neurons with randomized and equalized anatomical location (CA1 or SUB). Shuffling was performed on data in the open field (left column) or the spatial task (right column). The red bar indicates the difference of decoding error between the two neuronal populations recorded in the CA1 or the SUB, respectively. As we shuffled 1000 times, the p value of the red bar is smaller than 10e^−3^. (B) The blue histograms show the frequency of occurrence for differences in decoding error (left) and frequency of exactly decoded bins (FED) when decoding from populations of neurons with randomized task identity (OF or ST). Shuffling was performed on data from the CA1 (left column) or the SUB (right column). The black bar indicates the difference of decoding error between the two neuronal populations recorded during OF or ST, respectively. As we shuffled 1000 times, the p value of the red bar is smaller than 10e^−3^.

Table 1 Number of recorded neurons for every tetrode of the dataset.

In order to approximate the anatomical distribution of the recorded cells within CA1 and SUB, the respective regions were subdivided into proximal (p), middle (m) and distal (d) parts. The boundary between the respective subregions was approximated by dividing CA1 into three and SUB into two equally large subregions in the plane that the brains were sectioned.

